# On the determinants of residence times and dissociation mechanisms of complexes of interleukin-13 with its low and high affinity receptors

**DOI:** 10.64898/2026.08.13.743369

**Authors:** Nico Herb, Mislav Brajković, Giulia D’Arrigo, Daria B. Kokh, Rebecca C. Wade

**Affiliations:** Molecular and Cellular Modeling Group, Heidelberg Institute for Theoretical Studies, Schloss-Wolfsbrunnenweg 35, 69118 Heidelberg, Germany; Heidelberg Biosciences International Graduate School (HBIGS), Heidelberg University, Im Neuenheimer Feld 501, 69120 Heidelberg; Faculty of Biosciences, Heidelberg University, Im Neuenheimer Feld 234, 69120 Heidelberg; Faculty of Engineering Sciences, Heidelberg University, 69120 Heidelberg; Center for Molecular Biology of Heidelberg University (ZMBH), DKFZ-ZMBH Alliance, Im Neuenheimer Feld 329, 69120 Heidelberg, Germany; Interdisciplinary Center for Scientific Computing (IWR), Heidelberg University, Im Neuenheimer Feld 205, 69120 Heidelberg, Germany

**Keywords:** Interleukin-13, cytokine receptor, residence time, dissociation rate, molecular dynamics simulation

## Abstract

Interleukin-13 (IL-13) is an immunomodulatory cell signaling cytokine that has been implicated in neurodegenerative disease and chronic inflammation. IL-13 binds to its low and high affinity receptors, IL-13 receptor α1 (IL-13Rα1) and IL-13 receptor α2 (IL-13Rα2), respectively, with residence times that vary accordingly. As the binding kinetics of the cytokine-receptor complexes influence cellular responses, we employed the molecular dynamics (MD) simulation-based *τ-*random acceleration molecular dynamics method (*τ*RAMD) to compute relative residence times for wild-type (WT) IL-13 and 19 IL-13 mutants to the two receptors. Comparison with experimental kinetic data shows that the *τ*RAMD computations capture the trends in residence times. Analysis of simulated dissociation trajectories of the cytokine-receptor complexes reveals two distinct dissociation pathways of IL-13 from each of the receptors. This study thus pinpoints key determinants of the interaction of IL-13 with its receptors which could be targeted for therapeutic design.

**Statement of Significance:** Cytokines are regulatory proteins that bind to cell surface receptors and thereby send signals to the cellular interior. Interleukin-13 (IL-13) is a cytokine that has a low and a high affinity receptor. It has important physiological roles, and its deregulation is involved in diseases such as atopic dermatitis and asthma. We employed a molecular dynamics simulation-based method to compute the effects of changes in the sequence of IL-13 on the lifetimes of complexes of IL-13 and its receptors. Comparison with experiments supports the validity of the computational approach and analysis of the simulations reveals two distinct ways in which IL-13 dissociates from each receptor. These results thus provide a map for targeting IL-13 – receptor interactions for the design of therapeutics.

## Introduction

Interleukins are secreted cytokine proteins that bind to specific receptors on cellular surfaces (**1, 2**). The formation of an interleukin-receptor complex triggers a downstream signaling event in which intracellular signal transducers and activators of transcription (STATs) are phosphorylated by Janus tyrosine kinases (JAKs) that associate with the intracellular part of the receptor. Subsequently, the STATs form dimers and migrate into the nucleus to activate the transcription of target genes, see **Figure 1**.

**Figure 1:**
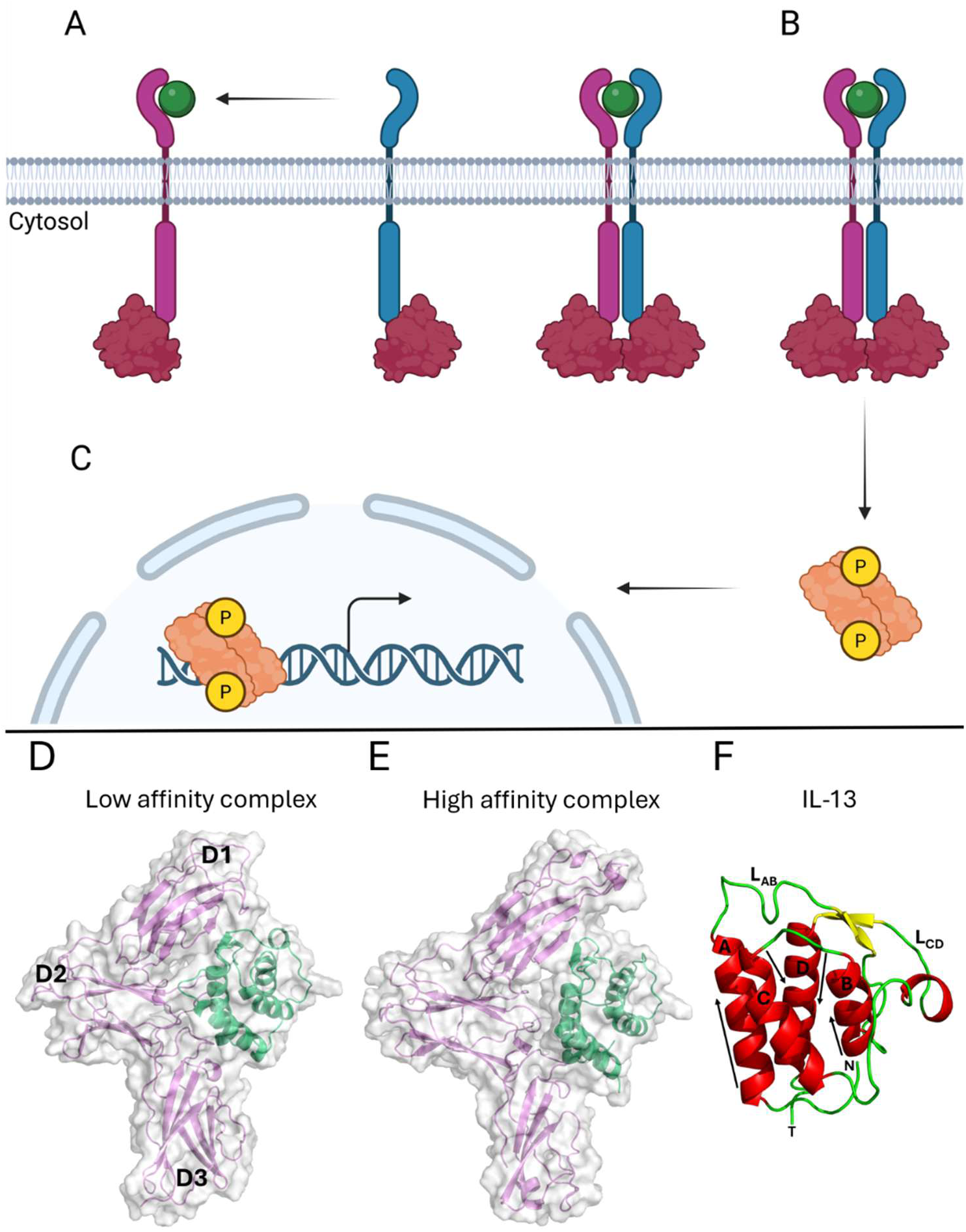
Schematic overview of the interleukin-dependent JAK/STAT signaling pathway and the three-dimensional structures of IL-13 and the binary low and high affinity IL-13-receptor complexes. Formation of a complex between IL-13 (green sphere) and the interleukin receptors (magenta and blue) occurs in a sequential manner **(A)** and is followed by **(B)** JAK-dependent (red) phosphorylation (yellow spheres) and dimerization of STATs (orange), which then **(C**) migrate into the nucleus to activate gene transcription (curved arrow). Crystal structures of the binary complexes of IL-13 (green) and the extracellular domains of its receptors (magenta) are shown for the low affinity IL-13Rα1 (PDB (**www.rcsb.org**) ID: 5E4E (**3**)**) (D)** and high affinity IL-13Rα2 (PDB ID: 3LB6 (**4**)**) (E)** receptors in ribbon and surface representation with their upper, middle and lower domains labelled D1, D2 and D3, respectively. The secondary structure elements of IL-13 are shown in **(F)**. The four helices, A-D, adopt an up-up-down-down topology as depicted by the black arrows. Two long loops connect helices A and B (L_AB_), and helices C and D (L_CD_). The N- and C-termini are indicated by the letters N and T, respectively. Helices are shown in red, an antiparallel β-sheet in yellow, and loops in green. All structures were visualized using the PyMOL Molecular Graphics System, Version 3 (**5**)**. (A)-(C)** were adapted from **Saxton et al.** (**2**) and created in BioRender. Wade, R. (2026) https://BioRender.com/cuslhqe.

An immune response is triggered upon communication between antigen-presenting cells and naive T cells. The subsequent differentiation of naive T cells into different types of T-helper cells results in the production of a specific set of interleukins, depending on the type of T-helper cell. Here, we focus on IL-13, which is produced by T-helper type II cells (**6**). It is a small protein consisting of four left-handed α-helices in an up-up-down-down topology (**7, 8**). Once secreted, IL-13 can bind to IL-13Rα1 and interleukin-4 receptor alpha (IL-4Rα) to form a ternary complex (known as a ternary type II complex, see **Figure 1A**) and IL-13Rα2 to form a binary complex (**6, 9, 10, 11**). Both complexes have been characterized extensively by **LaPorte et al.** (**12**)**, Lupardus et al.** (**4**) and **Moraga et al.** (**3**) and their structures are depicted in **Figure 1D** and **E**.

IL-13 first binds to IL-13Rα1 and subsequently, IL-4Rα is recruited to form a biologically active ternary complex (**Figure 1A**) (**12–14**). Several studies have demonstrated that IL-13 displays a nanomolar affinity to IL-13Rα1 (**3, 4, 12–17**). The reported affinities of the binary complex between IL-13 and IL-13Rα2 are in the nanomolar to picomolar regime (**13, 15, 18–23**). An extraordinarily high affinity in the femtomolar regime has been reported by **Lupardus et al.** (**4**). These studies showed that IL-13 displays a higher affinity towards IL-13Rα2 than to IL-13Rα1. As a result, IL-13 in complex with IL-13Rα1 is referred to as the “low affinity complex” whereas IL-13 in complex with IL-13Rα2 is referred to as the “high affinity complex”.

In addition to the higher affinity, it has been found that IL-13Rα2 can downregulate IL-13 mediated signaling since IL-13Rα2-deficient mice had increased fibrosis and the application of an engineered soluble form of IL-13Rα2 decreased fibrosis in IL-13Rα2-deficient mice (**24**). Furthermore, IL-13Rα2 did not activate STAT6 in the presence of IL-13, which contrasts with IL-13Rα1/IL-4Rα-dependent STAT6 activation (**21**), and, consistently, it has a short cytoplasmic region lacking signaling motifs (**20**). Therefore, IL-13Rα2 has been regarded as a decoy receptor. However, recent studies show that IL-13Rα2 does not function solely as a decoy receptor (**10**). There is evidence that, in the presence of IL-13, a full-length IL-13Rα2 is necessary for the production of transforming growth factor (TGF) β1, which contributes to fibrosis (**25**). Additionally, IL-13 is not the only ligand binding to IL-13Rα2. The chitinase-like protein (Chi3l1/YKL-40) can also bind to IL-13Rα2, which results in the phosphorylation of protein kinase B (PKB/AKT) and the mitogen activated protein kinase (MAPK) ERK (extracellular-signal related kinase), as well as in the translocation of β-catenin into the nucleus, contributing among other effects to inflammasome and oxidant injury inhibition (**26**).

Parkinson’s disease and Alzheimer’s disease are two of the most prominent forms of neurodegenerative diseases whose occurrence increases with age (**27**). It has been predicted that, worldwide, by the year 2040, 12.9 – 14.2 million individuals will be affected by Parkinson’s disease (**28**), and by 2050, over 106 million people will have Alzheimer’s disease (**29**). Both neurodegenerative diseases are associated with an elevated expression of IL-4Rα and IL-13Rα1 (**10**). For example, it has been demonstrated using mouse models that dopaminergic neurons in the substantia nigra pars compacta region of the brain express IL-13Rα1, which enhances the susceptibility of the respective neurons to oxidative stress resulting in their loss (**30**). However, the loss of dopaminergic neurons under chronic stress conditions is postponed in the absence of IL-13Rα1 in mouse models (**31**). These studies provide evidence that, at least under stress conditions, IL-13Rα1 is involved in the loss of neurons, which is one of the hallmarks of neurodegenerative diseases (**32**). IL-13 is naturally expressed in the central nervous system, but there is evidence that it can have both beneficial and detrimental effects on neurons (**11**). By using mouse and rat models, the localization of IL-13 has been demonstrated to be pre-synaptic while IL-13Rα1 is mainly post-synaptic (**33**). Mouse models have shown that IL-13 has beneficial effects after traumatic brain injury (**34**) whereas in the immune system, IL-13 is associated with asthma and atopic dermatitis (**9**) and humanized monoclonal antibodies that act as IL-13 antagonists by interfering with IL-13 – receptor binding are used in the clinic for treating these conditions (see e.g. https://www.ema.europa.eu/en/medicines/human/EPAR/ebglyss). There is also evidence for the involvement of IL-13Rα2 in cancer, and its (over-)expression has been observed in several cancers including glioblastoma, melanoma, ovarian, pancreatic, breast, colorectal, lung and prostate cancers (**35**).

The dissociation rate *k*_off_, is a binding kinetic parameter that describes the rate at which a receptor-ligand-complex separates into its two components. Its inverse, the residence time τ, is an important pharmacological measure that describes how long a ligand and a receptor stay bound to each other (**36–38**). Studies have shown a positive correlation between the protein-agonist residence time and agonist efficacy, with, for example, drugs with higher residence times increasing the survival rate of bacteria-infected mice (**39, 40**). However, high residence times can also induce unwanted side effects. An example is given by the drug roxifiban, which induces a conformational change of its target protein that is located on platelets but the altered conformation of the target protein is recognized by the immune system, resulting in platelet depletion (**41–43**). The dissociation rate or residence time is thus an important parameter to consider in the drug discovery process but is challenging to predict with conventional MD simulations because the clinically relevant time scales are not computationally accessible (**44**).

We here employ τ-RAMD, a procedure to compute relative residence times. It employs RAMD simulations in which, in addition to the standard force field, an additional randomly oriented force is adaptively applied to the center of mass (COM) of the ligand of a receptor-ligand complex in order to accelerate the unbinding event so that it can be observed in trajectories on the nanosecond time scale (**45, 46**). While the calculated residence times are much shorter, they can correlate with experimental values if the RAMD simulations capture the features responsible for the differences in residence time within a set of complexes, e.g. of a protein with a set of small molecules or of a set of protein mutants (**45, 47**). For example, calculated relative residence times compared well with experimental ones for 70 inhibitors that bind to heat shock protein (HSP) 90 (**45**), for compounds that bind to G-protein coupled receptors (**48**), and for mutants of the barnase-barstar protein-protein complex (**47**). To understand how ligands dissociate from their respective receptors and what type of interactions occur during the dissociation process that might influence the residence time, we used the Molecular Dynamics-Interaction Fingerprint (MD-IFP) tool (**49**), which was developed to study dissociation routes of protein-small molecule complexes (**49**) and protein-protein complexes (**47**), to analyze the RAMD trajectories.

In this work, *τ*RAMD was utilized to compute the residence times of IL-13 variants in complex with IL-13Rα1 or IL-13Rα2 to assess the performance of τ-RAMD on this important representative of cytokine-receptor protein-protein complexes and to gain insights into the determinants of their residence times. Three datasets were considered: Complexes with the high-affinity IL-13Rα2 and WT IL-13 and IL-13 mutants carrying a single point mutation to alanine in the binding interface (**4**); complexes of the low-affinity IL-13Rα1 and the same set of IL-13 variants plus additional IL-13 variants (**4**); complexes of the low-affinity IL-13Rα1 and WT IL-13 and several IL-13 variants with multiple mutations (**3**), see **Table 1 and Table S1**. Most of the mutations are localized at the protein-protein interface, see **Figure 2**. A plot of *k*_off_ versus *K_D_* values measured by surface plasmon resonance shows that they are highly correlated with a positive correlation of R=0.95 (R^2^=0.91). For two out of three datasets studied, the trend in residence times was captured and overall, a good agreement between computed and experimental residence times was achieved using *τ*RAMD. Analysis of the simulated dissociation trajectories revealed two dissociation pathways for the low and high affinity complexes, respectively, and enabled identification of important interactions between IL-13 and its receptors that can be exploited for therapeutic design.

**Figure 2:**
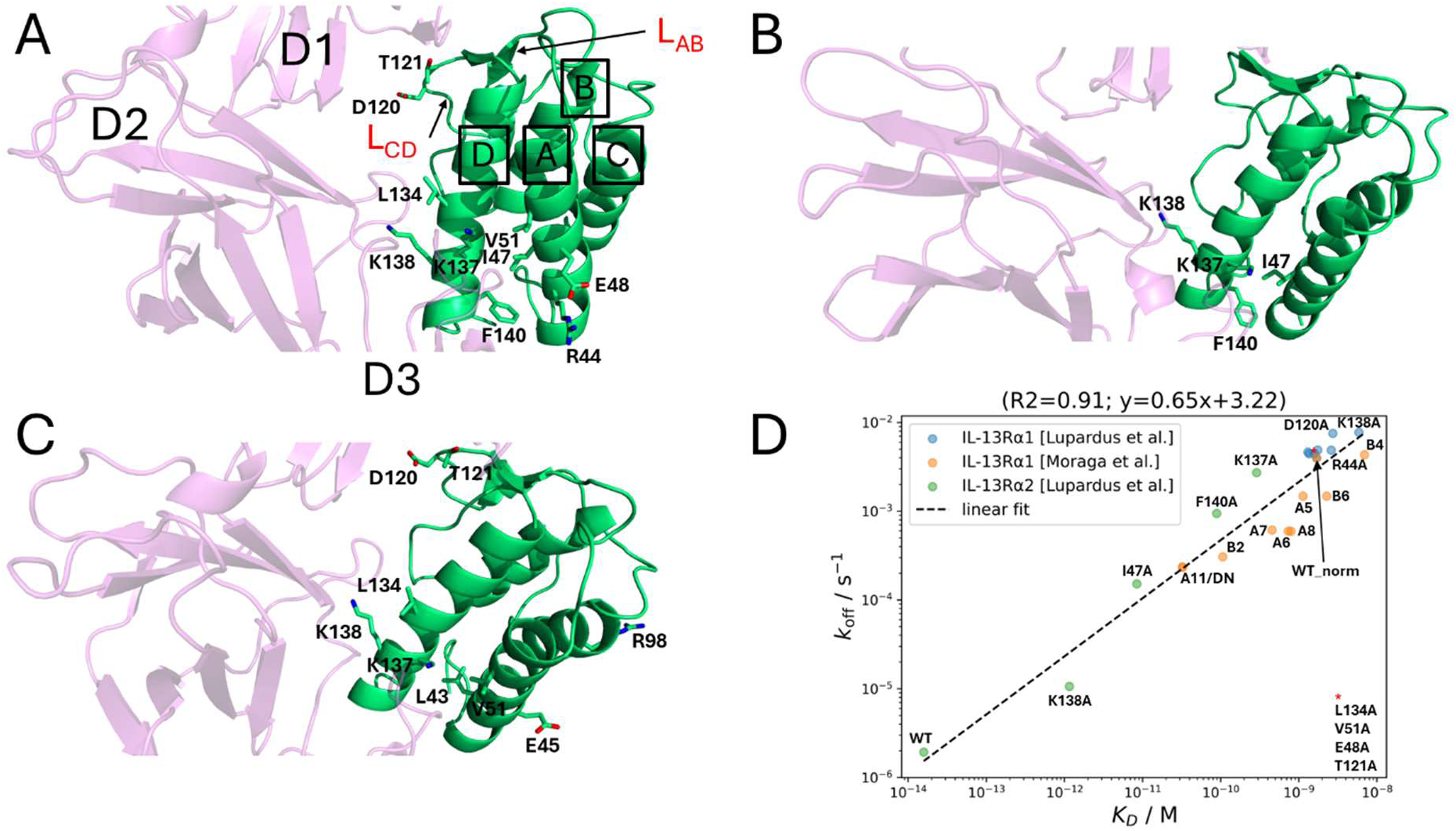
Structural localization of the IL-13 mutations studied and the corresponding measured binding parameters. The positions of mutations in IL-13 are shown in the complex with IL-13Rα1 **(A)** and with IL-13Rα2 **(B)** (**4**) and, as a representative mutant in the measurements by (**3**), for DN IL-13 in complex with IL-13Rα1 **(C)**. The three domains of IL-13Rα1 are labeled D1-D3. IL-13 helices are labelled A-D. Loops L_AB_ and L_CD_ are labeled in red and mutated residues (listed in **Table 1**) are shown as sticks colored by atom type. Experimentally measured *k*_off_-rates are plotted versus measured *K_D_* values in **(D)**. A linear regression in logarithmic space is shown by the dashed line. Error bars are not available. The dataset from **Moraga et al.** (**3**) (orange data points) was normalized such that the datapoint of WT IL-13 coincides with WT IL-13 of the dataset for the complex with IL-13Rα1 from **Lupardus et al.** (**4**) (blue data points). This data point is labelled WT_norm. The red asterisk represents a cluster of mutants with similar experimental values in **Lupardus et al.** (**4**).

**Table 1:** IL-13 variants modelled in binary complexes with IL-13Rα1 or IL-13Rα2. The measured binding kinetic parameters of the IL-13 variants are reported in the two references indicated. The residue numbering corresponds to the respective UniProt sequence. The measured data are given in **Table S1** along with the residue numbering in the respective references.

| Reference |  | Mutations | Name |
| --- | --- | --- | --- |
| Lupardus et al. (4) | IL-13Rα2 | -----<br>F140A<br>I47A<br>K137A<br>K138A | WT<br>F140A<br>I47A<br>K137A<br>K138A |
| | IL-13Rα1 | -----<br>F140A<br>I47A<br>K137A<br>K138A<br>R44A<br>E48A<br>V51A<br>D120A<br>T121A<br>L134A | WT<br>F140A<br>I47A<br>K137A<br>K138A<br>R44A<br>E48A<br>V51A<br>D120A<br>T121A<br>L134A |
| Moraga et al. (3) | IL-13Rα1 | R44I, V51I, R119K, D120G, T121S, L134H, K137R, K138A, F140M<br>L43I, V51L, R119M, D120K, T121S, L134H, K137R, K138A<br>L43V, V51I, D120S, T121S, L134F, K137R, K138T<br>V51I, R119T, D120G, T121S, L134Y, K137R, K138A<br>L43I, V51I, D120G, T121S, L134H, K137R, K138A<br>L43I, V51I, D120S, T121S, K122R, L134H, K137R, K138T<br>R44S, V51I, R119K, D120G, T121S, K122M, L134Y, K137R, K138T<br>L43T, V51I, D120G, T121S, L134Y, K137R, K138T<br>L43V, E45A, V51I, R98D, D120S, T121S, L134F, K137R, K138T | A5<br>A6<br>A11<br>B2<br>A7<br>A8<br>B4<br>B6<br>DN |

## Materials and Methods

### Experimental data

Experimental values of dissociation rates measured by surface plasmon resonance were taken from two studies (**3, 4**). The names of all IL-13 variants studied in complex with IL-13Rα2 or IL-13Rα1 are given in **Table 1**. The UniProt numbering (**50**) is used throughout this study (IL-13: P35225; IL-13Rα2: Q14627; IL-13Rα1: P78552).

Two crystal structures were extracted from the PDB for modelling the protein structures: the binary IL-13 : IL-13Rα2 complex (PDB ID: 3LB6) (**4**) and the ternary type II IL-13 : IL-13Rα1 : IL-4Rα complex (PDB ID: 5E4E) (**3**).

### Protein structure modeling

Based on the crystal structures, two models were prepared consisting of WT IL-13 in complex with the three extracellular domains (D1-D3) of either IL-13Rα2 or IL-13Rα1 by filling gaps, adding missing sidechains, and mutating residues to the WT sequence where necessary using SWISS-MODEL (**51**) and Maestro (**52**).

For the high affinity complex with IL-13Rα2, IL-13 was modelled from residues 38 to 146 as residues 1-37 were missing from the crystal structure and the engineered mutation Q144R (which is not directly located at the protein-protein interface) in the PDB file was kept to match the sequence of IL-13 in the crystal structure of the high affinity complex. IL-13Rα2 consisted of residues 31 to 328, corresponding to those in the crystal structure. D279 and E291 were manually deprotonated where necessary. A missing oxygen atom was added to the C-terminal carboxy group of N146 of IL-13.

For the low affinity complex with IL-13Rα1, IL-13 was modelled from residues 34 to 146 as residues 1-33 were missing from the crystal structure, and the 8 mutations in IL-13 were reverted to match the UniProt sequence (V43L, I51V, R72L, S120D, S121T, F134L, R137K, T138K). IL-13Rα1 consisted of residues 32 to 340, corresponding to those in the crystal structure.

It should be noted that the proteins employed for the experimental dissociation rate measurements differed as follows:

- For surface plasmon resonance (SPR) measurements, low and high affinity complexes were expressed in glycosylated form. Crystallization was conducted so as to prevent glycosylation. However, the crystal structure of the high affinity complex shows a glycosylation site but this is not in proximity to the protein-protein interface.
- Moraga et al. (**3**) used untruncated human IL-13 and IL-13Rα1 including residues G23-Q332.
- Lupardus et al. (**4**) used constructs including IL-13 residues G35-N146 and IL-13Rα2 residues E29-W331.

With respect to *τ*RAMD, it is likely that these differences in sequence have only a minor effect on the calculation of the relative residence time and will most likely not change the trends in the dissociation pathways since the N- and C-termini are not located in the proximity of the interface between the proteins.

These models of WT IL-13 in complex with IL-13Rα2 or IL-13Rα1 served as templates for introducing mutations in IL-13 using Maestro versions 2021-4 and 2025-1 (**52**). All IL-13 variants for which structural models were made are listed in **Table 1**.

### System preparation

The procedures for system preparation, energy minimization, heating and equilibration with AMBER together with re-equilibration and the τ-RAMD production run with GROMACS, were adapted from **D’Arrigo et al.** (**47**). Simulation conditions were chosen to approximately correspond to the experimental conditions for measuring binding kinetics: Moraga et al. (**3**) purified IL-13 and IL-13Rα1 in 10 mM HEPES (pH 7.2) and 150 mM NaCl, whereas Lupardus et al. (**4**) purified IL-13, IL-13Rα2 and IL-13Rα1 for their biophysical measurements in 5 mM HEPES (pH 7.0) and 150 mM NaCl.

All of the complexes of the IL-13 variants with IL-13Rα2 or IL-13Rα1 were preprocessed using the Schrödinger preprocessing tool (**53, 54**) and protonated at pH 7.4 using PROPKA (**55, 56**) in Maestro versions 2025-1 and 2021-4 (**52**), respectively. The N-and C-termini of the modelled proteins were assigned their zwitterionic form and histidine residues 106 of IL-13 (in the high affinity complex) and 106 of IL-13Rα2 were doubly protonated.

Next, tLEaP from AMBER 18 (**57**) was used to place each model into a truncated octahedral periodic simulation box in which the minimal distance between the protein complex and the box edges was 25 Å to allow for dissociation. The simulation box was solvated with TIP3P water molecules (**58**) and a salt concentration of 150 mM NaCl was used to mimic experimental conditions (**3, 4**) and to neutralize the systems to a net charge of zero. All systems were prepared with the AMBER ff14SB (**59**) force field.

### Energy minimization, heating and initial equilibration

AMBER 20 (**60**) was used to carry out energy minimization, heating and initial equilibration. All prepared systems, consisting of a complex between different IL-13 variants and IL-13Rα2 or IL-13Rα1, were energy-minimized in four minimization runs using the steepest descent algorithm. The method of minimization was changed to conjugate gradient after 500, 100, 500, and 500 steps, respectively, for a total of 1000, 1400, 1000, and 1000 steps, respectively. During the first three energy minimizations, harmonic restraints were applied to all heavy atoms of both proteins with a force constant of 500, 100, or 5 kcal mol^-1^ Å^-2^, respectively. The last minimization was carried out without restraints. After energy minimization, all systems were heated to 300 K in a total of 200 ps while harmonically restraining all heavy atoms of the proteins with a force constant of 50 kcal mol^-1^Å^-2^ in the NVT ensemble using the Langevin thermostat. Next, all systems were equilibrated for 1 ns in the NPT ensemble keeping the temperature constant at 300 K using a Langevin thermostat and the pressure constant at 1 bar using a Berendsen barostat while restraining all heavy atoms of both proteins with a force constant of 50 kcal mol^-1^ Å^-2^. The equilibration was then repeated for 1 ns without restraining the systems. With respect to non-bonded interactions, a cutoff distance of 10 Å was used for energy minimization, heating and equilibration. Furthermore, periodic boundary conditions were applied to the systems together with the Particle Mesh Ewald method. For the equilibrations, SHAKE (**61**) was used to constrain bonds involving hydrogen atoms and the time step was set to 2 fs.

### Equilibration using GROMACS

Using the coordinate files generated at the end of the equilibration with AMBER 20 (**60**) and the topology file from tLEaP (AMBER 18, (**57**)), cpptraj (AMBER 24/22 (**62, 63**), respectively) and ParmEd (**64**) were used to generate the GROMACS topology and structure files for each system. These were used to perform a re-equilibration in a version of GROMACS 2020.5 (**65**) containing the RAMD 2.0 implementation (**49**) (GROMACS-RAMD version 2020.5-2.0). The first part of this equilibration was carried out in the NVT ensemble for a total of 10 ns using the Berendsen thermostat at 300 K with velocity generation based on the Maxwell-Boltzmann distribution at 300 K. The second part of the equilibration was carried out in the NPT ensemble using a Nosé-Hoover thermostat at 300 K and a Parrinello-Rahman barostat at 1 bar for a total of 20 ns. The second part consisted of six independent equilibrations of 20 ns each, each run with the same input coordinates and velocities, to generate six different starting points for the subsequent RAMD simulations.

### RAMD simulations

In total, 90 RAMD simulations were run for each system, i.e. 15 trajectories starting from each of the 6 replicas obtained after the second equilibration with GROMACS. The RAMD simulations were performed in the NPT ensemble with a maximum trajectory length of 40 ns. During the simulations, a randomly oriented force of constant magnitude of 1100 kJ mol^-1^nm^-1^ (∼ 26 kcal mol^-1^ Å^-1^) was applied to the IL-13 COM. Every 100 fs, it was evaluated whether the distance travelled by IL-13 exceeded a threshold distance of 0.025 Å. If not, the direction of the force was altered randomly, otherwise the simulation was continued without changing the force direction. When the distance between the IL-13 COM and the receptor COM exceeded 70 Å, the trajectory was stopped. If the trajectory was stopped prior to dissociation due to reaching the 40 ns time limit, then the respective RAMD trajectory was rerun. For some trajectories of the high affinity complexes, the maximum trajectory length was increased from 40 ns to 80 ns. If the standard deviation of the calculated relative residence time exceeded 50%, all trajectories were re-run with an additional five RAMD simulations per replica. Recomputation of coordinates to account for periodic boundary conditions was necessary prior to any of the subsequent post-processing analysis.

### Calculation of rRAMD residence times

The relative residence times of the complexes were calculated according to the original τRAMD publication (**45**). Once the distance between receptor COM and ligand COM exceeds a user-defined threshold distance, the trajectory is terminated and the length of the trajectory recorded. The RAMD residence time τ is calculated as the mean trajectory length from the set of replicas, each consisting of a set of independent trajectories from which a replica residence time is calculated. The simulation time after which 50% of all trajectories of one replica show a dissociation event is defined as the replica residence time. The distribution of the residence time is assessed by performing a bootstrapping for all trajectories of each replica (**45**). Here, bootstrapping was performed by randomly selecting 80% of all simulation times that belong to one replica and calculating the median. In total, this procedure was repeated 50 000 times. Then, an average of all 50 000 bootstrapped medians was calculated and this was defined as the residence time of a replica r_r_ together with its standard deviation *sd*_r_. The procedure was applied to all replicas and the calculated residence time was then defined as the mean of all replica residence times r_c_with the respective standard deviation *sd*_c_. The residence time distribution was assessed using a Kolmogorov-Smirnov test. A detailed explanation of the residence time calculation and the statistical analysis is provided by **Kokh et al.** (**49**). As an example, the calculation and statistical evaluation of the residence time for the complex of WT IL-13 and IL-13Rα1 is shown in **Figure S1**.

In addition to the stopping criterion based on interleukin COM to receptor COM distance (COM-COM), four criteria introduced by **D’Arrigo et al.** (**47**) were also utilized to calculate the residence time: “by residue first”, “by residue last”, “few contacts first”, and “many contacts last”. The “by residue first” criterion considers all frames of a trajectory up to first frame in which the average distance between residue-residue contacts exceeds a threshold distance of 5.5 Å and the “by residue last” criterion considers all frames up to the very last fame in which the average distance between residue-residue contacts was less than 5.5 Å. The “few contacts first” criterion considers all frames of a trajectory up to the first frame in which the number of residue-residue contacts falls below a threshold of 50% and the “many contacts last” criterion considers all frames up to the very last frame in which the number of residue-residue contacts was higher than 50% (**47**). Note that the COM-COM criterion was used in order to terminate the RAMD trajectories, whereas residence times according to the other criteria were computed only for a specific range of frames during the post-analysis of the RAMD trajectories generated with the COM-COM criterion. Again, bootstrapping was performed for the times extracted according to the different criteria.

### Analysis of the dissociation trajectories

MD-IFP analysis (**49**) was performed for all complexes as described by **D’Arrigo et al.** (**47**). Protein contacts were analyzed for a defined range of frames from all RAMD trajectories of a system. The range of frames extended from that at which the “by residue first” criterion was satisfied up to the last frame of a dissociation trajectory. In addition, to sample the bound state, 200 frames prior to this frame range were considered. For each frame analyzed, a matrix was built containing all residue-residue distances between the two proteins that were within 15 Å. For each pair of residues, the distance was defined as the smallest difference in the distance between the centers of mass of predefined sets of residue atoms. In addition, for each analyzed frame, the difference in the distance between the centers of mass of the two proteins (ΔCOM) and the root mean square deviation (RMSD) with respect to the original position of the protein to which the force is applied were saved.

The Jaccard distance was used to assess the dissimilarity between RAMD frames and then clusters were generated using the *k-means* algorithm (**66**). Based on elbow and silhouette analysis, the number of clusters was chosen to be five to capture the main features of the dissociation trajectories for all variants and both complexes.

Frames with fewer than three contacts were subject to hierarchical clustering and then displayed in heatmaps. In this part of the trajectory, where the two proteins are almost completely separated from each other, it is easier to distinguish between different dissociation pathways, especially when the population of alternative pathways is low.

The PyMOL Molecular Graphics System, Version 3 (**5**) was used for all protein structure visualization. Movies of trajectories were generated with VMD, version 1.9.4 **(67;** http://www.ks.uiuc.edu/Research/vmd/)

## Results

### Computed residence times capture the trends in the experimental residence times

The computed τRAMD residence times are plotted against the experimentally determined values in **Figure 3** and **Figures S2**, **S3** and **S4**. All experimentally determined *k*_off_-rates and the corresponding residence times and affinities are given in **Table S1** whereas the calculated relative residence times for all datasets and criteria are given in **Table S2**.

**Figure 3:**
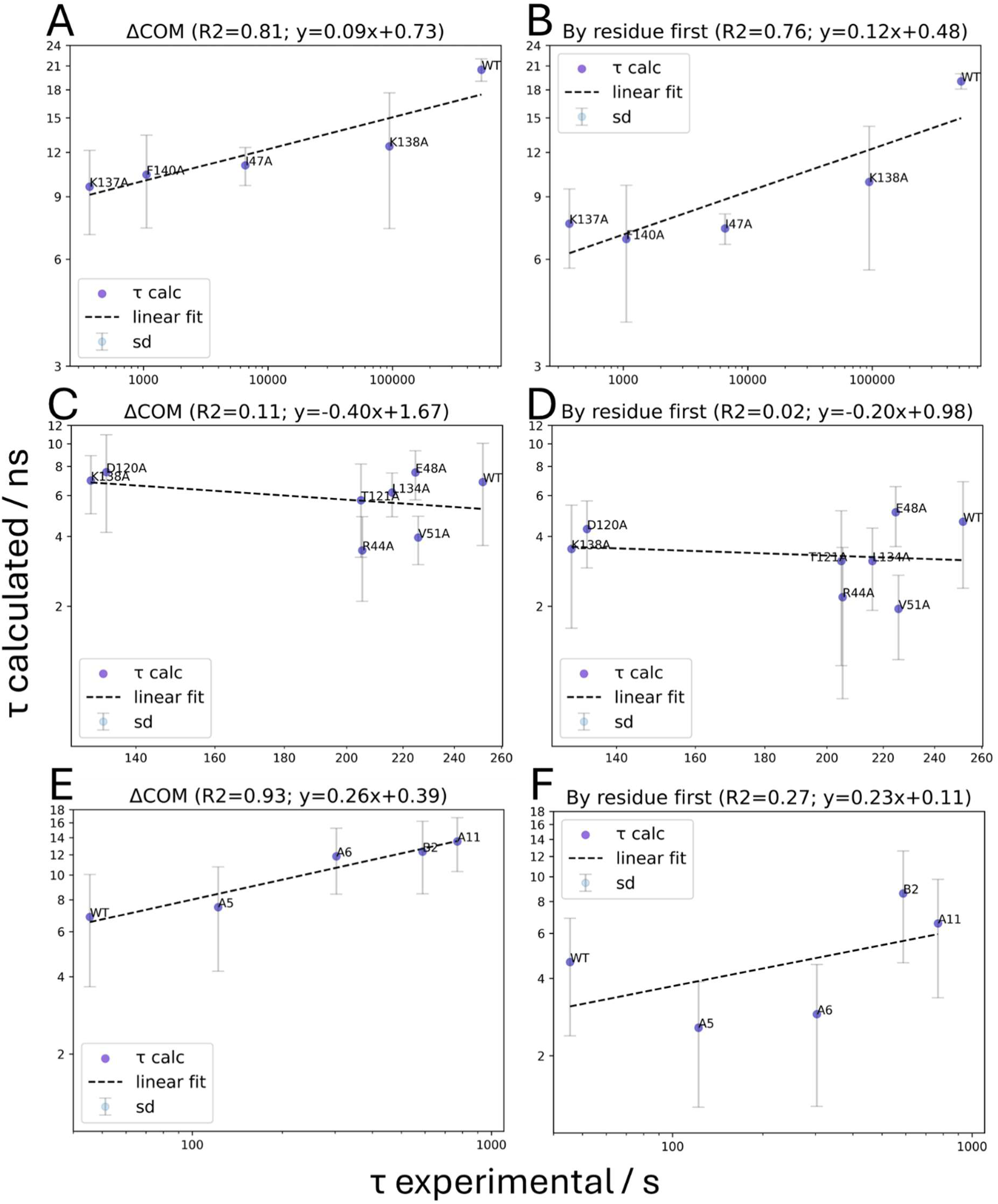
Comparison of calculated and experimental residence times for the three datasets. Relative residence times were calculated according to **(A,C,E)** the COM-COM and **(B,D,F)** the “by residue first” criteria (**47**) for IL-13 in complex with **(A,B)** IL-13Rα2 (**4**); **(C,D)** IL-13Rα1 for the IL-13 variants reported in Ref. (**4**); and **(E,F)** IL-13Rα1 for the IL-13 variants reported in Ref. (**3**). Linear regressions in logarithmic space are shown by the dashed lines. The data points give the mean value of six simulation replicas together with the corresponding standard deviation (see Methods for details).

The experimentally determined residence times of IL-13 in complex with IL-13Rα2 cover a wide range, extending from six minutes (K137A) to six days (WT) (**4**) (**Figures 3A** and **3B**). Using the COM-COM criterion, a high positive correlation of 0.9 with R^2^=0.81 was obtained between calculated and experimental residence times. The application of the “by residue first” criterion decreased the calculated residence times of all mutants and the WT and resulted in a slightly lower correlation with the experimental values (R^2^=0.76). The application of the remaining criteria (“by residue last”, “few contacts first” and “many contacts last”) resulted in only minor differences in the calculated residence times and no notable difference in the correlation, see **Figure S2**.

In contrast to IL-13Rα2, the residence times of IL-13 variants in complex with IL-13Rα1 determined experimentally by (**4**) are much shorter and cover a small range (**Figures 3C** and **3D**). The experimental residence times for WT and the R44A, E48A, V51A, T121A and L134A mutants are all rather similar, lying between 200 and 260 s, whereas IL-13 variants D120A and K138A have residence times of 133 and 130 s, respectively. Using the COM-COM criterion, a weak negative correlation of -0.33 (R^2^=0.11) was obtained. The application of the “by residue first” criterion again lowered the computed residence times but did not improve the correlation, yielding a weak negative correlation of -0.14 (R^2^=0.02). As for the high affinity complex, the application of the remaining criteria (“by residue last”, “few contacts first” and “many contacts last”) to the low affinity complex yielded similar results, see **Figure S3**.

A different set of IL-13 mutants and their residence times were published for the low affinity complex by **Moraga et al.** (**3**). In contrast to the dataset from **Lupardus et al.** (**4**), the residence times determined by **Moraga et al.** (**3**) are more distinct from each other and cover a wider range, from 42 to 769 s (**Figures 3E** and **3F**). Therefore, for the initial evaluation, we studied the WT and the A5, A6, A11 and B2 mutants, which have distinct experimental residence times. (The remaining mutants reported by **Moraga et al.** (**3**) - A7, A8, B4, B6 and DN - have similar or identical experimental residence times (**Table S1**)). The comparison between the experimental values for these five IL-13 variants and calculated residence times is shown in **Figures 3E** and **3F**.

Using the COM-COM criterion, a positive correlation of 0.96 (R^2^=0.93) was obtained. The application of the “by residue first” criterion gave decreased computed relative residence times with a markedly decreased correlation of 0.52 (R^2^=0.27). Again, the application of the remaining criteria (“by residue last”, “few contacts first” and “many contacts last”) resulted in only minor differences in the calculated residence times and no notable change in the correlation compared to the COM-COM criterion, see **Figure S4**.

So far, the three datasets were treated separately to examine intra-dataset variation. Next, to evaluate whether the calculated values distinguished between low and high affinity receptor binding, the dataset of the low affinity receptor from **Moraga et al.** (**3**) was normalized to that of **Lupardus et al.** (**4**) such that both experimentally determined residence times for WT IL-13 in complex with IL-13Rα1 coincide (the dataset of **Moraga et al.** (**3**) was shifted to longer residence times). Comparison of all computed values for the three datasets shows an overall good agreement between experimental and calculated residence times with a correlation of 0.73 (R^2^=0.54), see **Figure 4**.

**Figure 4:**
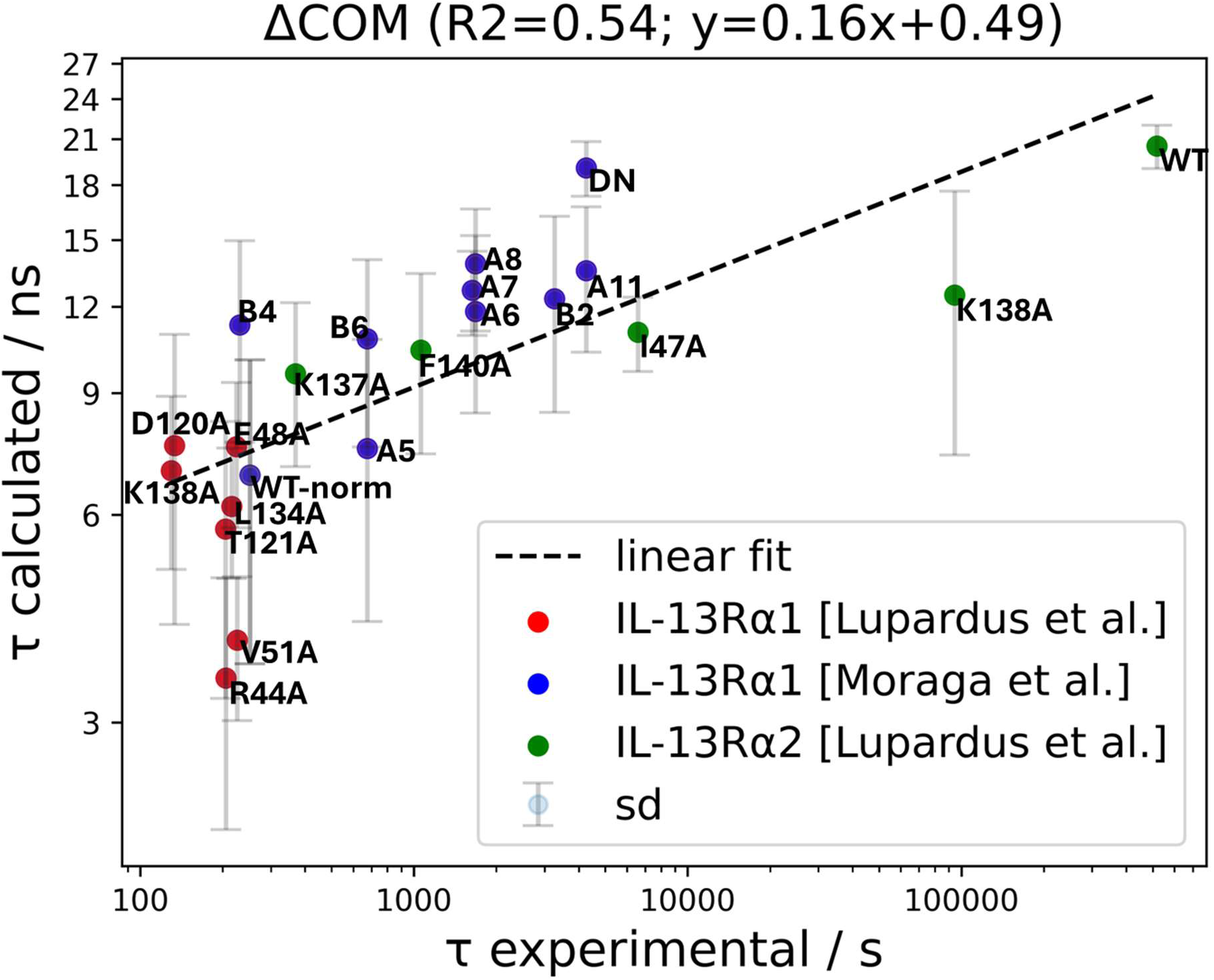
Comparison of calculated and experimental residence times for WT and all IL-13 variants in complex with IL-13Rα1 and IL-13Rα2. Residence times are computed according to the COM-COM criterion. A linear regression to the data points is shown by the dashed line. The data points give the mean value of six simulation replicas together with the corresponding standard deviation (see Methods section for details). All standard deviations are below 50% of the computed residence time value. Error bars are not available for the experimentally determined values. The dataset from **Moraga et al.** (**3**) (blue) was shifted to higher residence times so that the experimental residence times of WT IL-13 from the low affinity dataset of **Lupardus et al.** (**4**) (red) and the low affinity dataset of **Moraga et al.** (**3**) coincide. This is indicated by WT-norm (normalized).

In summary, we find that although small differences in the shorter residence times of the low affinity complexes measured by **Lupardus et al.** (**4**) cannot be reproduced, the overall trends in residence times can be captured by the τRAMD calculations for both low and high affinity receptor complexes.

### Analysis of dissociation trajectories reveals that both the low and the high affinity complexes have two dissociation pathways

To analyze the dissociation mechanisms of the IL-13 variants, MD-IFP analysis (**47**) was carried out for the RAMD trajectories of each IL-13 variant in complex with IL-13Rα2 or IL-13Rα1. Following silhouette and elbow analysis (**Figures S5 - S8**) to determine a suitable number of clusters to describe the dissociation trajectories, all analyzed frames for each complex were grouped into five clusters according to the residue-residue contact fingerprint content.

### Dissociation of the IL-13: IL-13Rα2 high affinity complex

The results of the analysis for WT IL-13 in complex with IL-13Rα2 are shown in **Figure 5**. Clusters 1 and 2 can be considered to represent the bound state since the increase in the distance between the COMs of the receptor and interleukin (ΔCOM) was only up to 1 Å on average (**Figure 5A**). Cluster 3 represents a loosening of the bound state with IL-13 displaced by about 3 Å. Cluster 4 represents a metastable state in which the interleukin has already moved slightly more than 10 Å from the receptor until it eventually dissociates to the unbound state, which is represented by cluster 5. In the bound state, IL-13 forms numerous contacts with IL-13Rα2 through a large interface (**Figure 5B**). D1 of the receptor is in contact with the loop regions L_AB_ and L_CD_ of IL-13 while D3 is in contact with the A-helix of the interleukin. Approximately half of all contacts are formed between the D-helix of IL-13 and domains D2 and D3 of the receptor, making the D-helix an important part of the interface. As the proteins start to dissociate, the predominant contacts between D1 (I84, I85, T86, K87 and N88) and L_CD_ (D120, T121) decrease in cluster 3 while the predominant contacts between D2 and D3 of the receptor (Y207, R268, I314, S317 and D318) and the D-helix of IL-13 (K137, K138 and F140) persist throughout clusters 1 to 3. In cluster 4, both these regions show a reduction in the number of contacts before the loss of all contacts in the unbound state, cluster 5. Thus, the interleukin tends to first lose contacts with D1 and then follow a dissociation route via D2 and D3. This dissociation pathway will be referred to as the reference pathway of the high affinity complex. Frames of a representative dissociation trajectory along this pathway are shown in **Figure 5E** and **Supplementary Movie M1**. It is worth mentioning that the trajectory shown does not include all residue-residue contacts shown in **Figure 5B**, since each RAMD trajectory individually contributes to the MD-IFP content in a different way. Therefore, the output of the MD-IFP analysis provides a contact-based overview of all trajectories.

**Figure 5:**
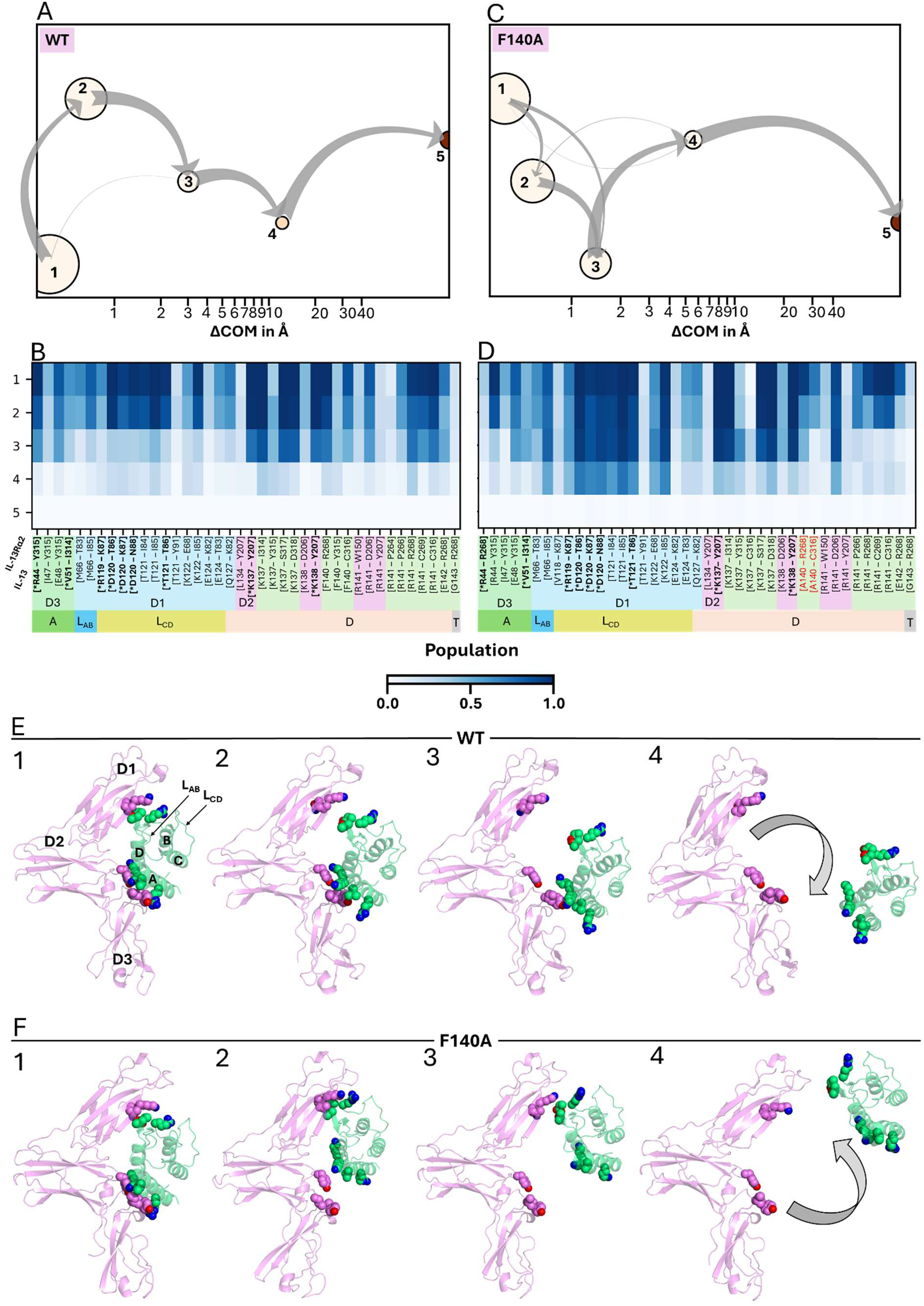
Analysis of RAMD trajectories of WT and F140A IL-13 dissociating from IL-13Rα2. All trajectories were analyzed to generate 5 structural clusters representing the predominant dissociation routes. These are shown in **(A, C)** where each circle represents one cluster (indicated by the number) positioned at its average increase in the distance between the centers of mass of the interleukin and the receptor (ΔCOM) on the logarithmic x-axis. The size of each cluster corresponds to its population, and the average RMSD value of each cluster is indicated by the color where darker indicates a higher RMSD value. The grey arrows indicate the flow between clusters. The residue-residue contacts made in the corresponding clusters (1-5 along the y-axis) are shown in the heat maps with darker color indicating higher population **(B, D)**. The residue-residue contacts were assigned to the specific parts of the two proteins: For the receptor: upper domain 1 (D1), light blue; middle domain 2 (D2), light purple; lower domain 3 (D3), light green. For the interleukin (lower bar): helix A (A), green; loop region between helix A and B (L_AB_), blue; loop region between helix C and D (L_CD_), yellow; helix D (D), beige; C-terminal flexible region (T), grey. Contacts formed by the F140A mutation are colored red. Representative frames from dissociation trajectories are displayed in **(E, F)** and corresponding **supplementary movies M1 and M2**. For both trajectories, frame 1 corresponds to the bound state. Frame 2 corresponds to a loosely bound state since IL-13 has lost contacts between L_AB_ and L_CD_ and D1, while still being associated with the receptor through D2 and D3 and helices A and D for WT IL-13. In contrast, the IL-13 F140A mutant has lost contacts between helices A and D and domains D2 and D3 while still being bound to D1 via L_AB_ and L_CD_. Frame 3 shows the progression of the dissociation for WT IL-13 where the interleukin starts to also lose contacts between D2 and D3 and helices A and D until it has completely dissociated from its receptor in the last frame 4, corresponding to the unbound state. IL-13 F140A starts to lose contacts between D1 and L_AB_ and L_CD_ until it has dissociated from its receptor in frame 4. IL-13 is colored green and IL-13Rα2 magenta. Residue contacts highlighted with an asterisk in **(B, D)**, given with IL-13 residue number (lower row) followed by IL-13Rα2 residue number, are depicted as spheres where oxygen atoms are colored red and nitrogen atoms are colored blue. The three domains of IL-13Rα2 are labeled D1, D2, and D3. Helices A, B, C, D and loops L_AB_ and L_CD_ are labeled on IL-13.

The analysis for the IL-13 F140A variant shows intermediate clusters 3 and 4 closer to the bound state than for WT IL-13 (**Figure 5C**). Comparison of the contact maps of this mutant and WT IL-13 (**Figure 5B** and **5D**) reveals a major difference in the persistence of the residue-residue contacts between D1 of the receptor and L_CD_ of the interleukin. While for the WT IL-13, these contacts are already lost in cluster 3, these contacts are maintained up to cluster 4 for the F140A mutant. Indeed, the contacts between D1 (I84, I85, T86, K87 and N88) and L_CD_ (D120 and T121) are longer lasting than the contacts between domains D2 and D3 and helix D, revealing an alternative pathway with a considerable population in comparison to the reference pathway.

Visual inspection of the RAMD trajectories revealed that the reference pathway for WT IL-13 in complex with IL-13Rα2 can also be observed for IL-13 F140A in complex with its high affinity receptor. However, a considerable number of trajectories show the alternative pathway in which the interleukin first loses contacts between D2 and D3 of the receptor and the A- and D-helices of the IL-13 F140A variant and then dissociates via D1, see **Figure 5F** and **Supplementary Movie M2**. The cluster representations and contact maps of all the IL-13 variants in complex with IL-13Rα2 are shown in **Figures S9** and **S10**, respectively. A comparison between the contact maps shows that the IL-13 I47A variant shows a similar contact pattern to F140A, indicating a considerable population of the alternative dissociation pathway. In contrast, the IL-13 K138A variant shows a similar contact pattern to WT IL-13, indicating that this mutant follows the same dissociation pattern as WT IL-13. Interestingly, IL-13 K137A shows an intermediate pattern in its contact map, indicating a more equal distribution of the reference and alternative dissociation pathways.

These findings are supported by hierarchically clustered fingerprints of the frames towards the end of the dissociation trajectories in which fewer than three contacts were present, see corresponding contact maps in **Figure S11**. The most prominent and last contacts in the reference egress pathway are between R141 of the interleukin and several residues of D2 and D3 in the receptor. On the alternative pathway, the most prominent and last contacts are between D120, T121 and K122 of the interleukin and E68, I85, K87 and E118 of the receptor.

### Dissociation of the IL-13: IL-13Rα1 low affinity complex

For WT IL-13, the clustering of each analyzed frame of the trajectories of the dissociation from IL-13Rα1 gave clusters 1-3 corresponding to the bound state, with cluster 3 being a loosened complex (**Figure 6A**). The average displacement of IL-13 from the receptor in the intermediate cluster 4 exceeded 10 Å. The corresponding contact map (**Figure 6B**) shows that all three domains of the receptor make contacts with various regions of IL-13 in the bound state. Specifically, IL-13 helices A and D, as well as the loops L_AB_ and L_CD_, are involved in contact formation. D1 forms numerous contacts with L_AB_, L_CD_ and the D-helix of IL-13. D2 is only in contact with the D-helix, and D3 is in contact with the A-helix, the D-helix and partially with the flexible part of the C-terminus. The D-helix forms approximately as many contacts with the receptor as the A-helix, L_AB_ and L_CD_ combined, thus being responsible for most of the contacts between the two proteins. Some of these contacts are maintained throughout clusters 1 to 3, though there is a reduction of contacts between D3 and the A-helix in cluster 3. The contacts between D1 and L_CD_ and between D1 and L_AB_ are maintained through to cluster 4. This indicates that IL-13 mainly first loses contacts between D3 and the A-helix, and D2/D3 and the D-helix, and then dissociates via the contacts maintained by D1 and L_AB_/L_CD_. In particular, the high occupancy contacts between D75, K76, K77, I78, A79, T104 and E106 of D1 and M66, T121, K122 and E124 of IL-13 tend to be the last contacts before dissociation. This dissociation pathway will be referred to as the reference pathway for the low affinity complex. Four frames of an exemplary trajectory for WT IL-13 are depicted in **Figure 6E** and the trajectory is shown in **Supplementary Movie M3**. The first frame represents the bound state (clusters 1 and 2). The second frame corresponds to a state in which the interleukin is loosely bound to its receptor (cluster 3). This is illustrated by the loss of numerous contacts between both proteins. In particular, while contacts are disrupted between D2 and D3 and helices A and D, contacts between D1 and L_AB_ and L_CD_ are maintained. The third frame shows the transition between the loosely bound state and the dissociated state in the last frame. Helices A and D rotate away from the receptor, and the interleukin slightly shifts sideways to the upper part of the receptor while still being bound via D1 and L_AB_ and L_CD_. The last frame in **Figure 6E** shows that the interleukin has lost the longest lasting contacts between D1 and L_AB_ and L_CD_, by “flipping” to the upper domain of the receptor from its bound position until it fully dissociates.

**Figure 6:**
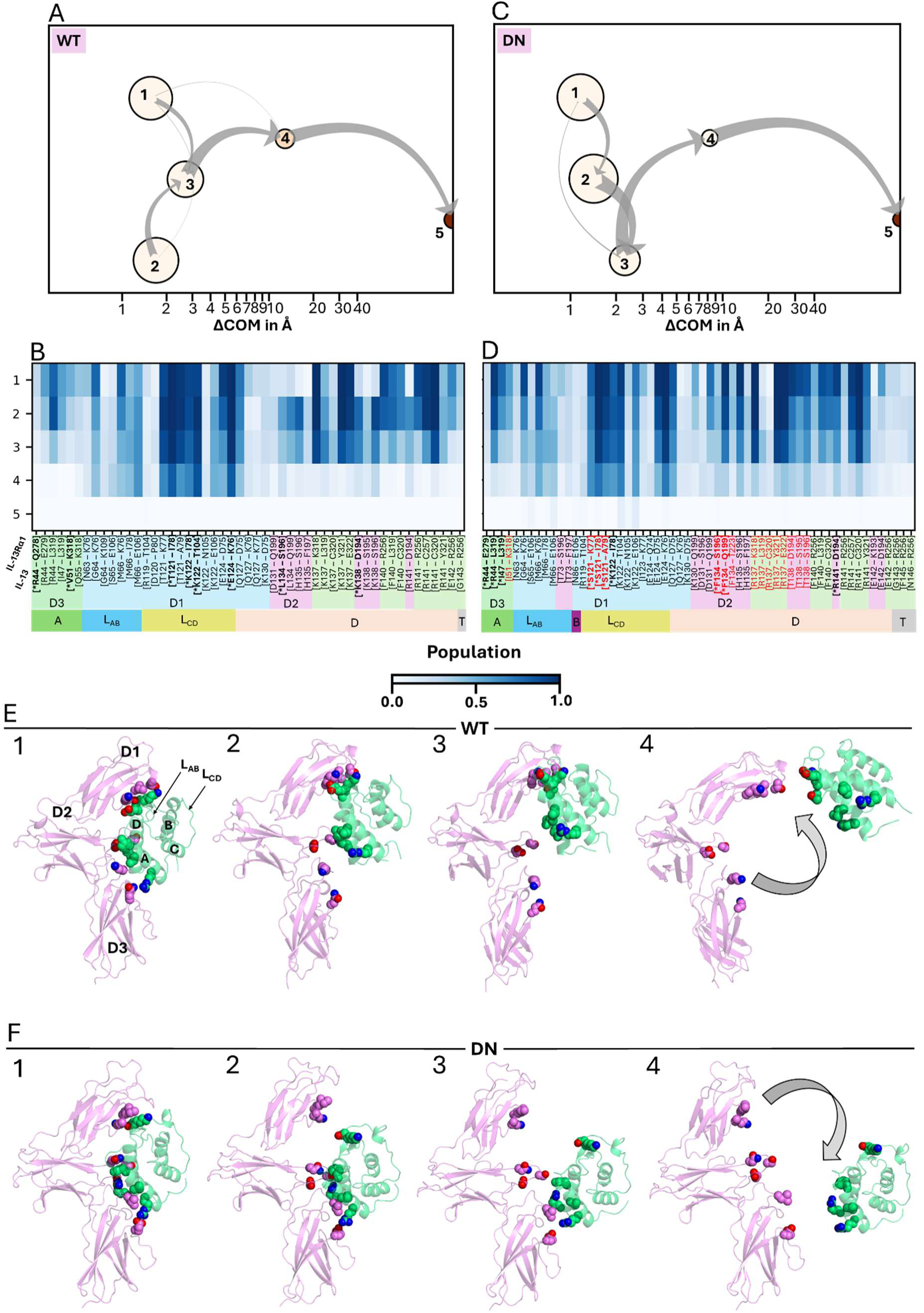
Analysis of trajectories of WT and DN IL-13 dissociating from IL-13Rα1. **(A, C)** Dissociation trajectories between clusters; **(B, D)** Residue-residue contacts in the clusters; Contacts in which mutations of DN IL-13 are present are colored red. **(E, F)** Frames of representative dissociation trajectories which are also shown in the corresponding **supplementary movies M3 and M4**. For details, see the legend for Figure 5.

A comparison between WT IL-13 and the two mutants with the highest experimental residence time out of all low affinity complexes under study, DN and A11, revealed no major differences in the cluster composition and mode of dissociation (see **Figure 6C** for DN). Clusters 1 to 3 resemble the bound state, with cluster 3 being a slightly loosened complex. Analogous to WT, cluster 4 represents a metastable state in which the interleukin is displaced by approximately 9 Å on average and cluster 5 corresponds to the dissociated state.

Besides some minor differences in the contact composition between the WT and the mutant complexes, the most intriguing difference is that, in comparison to the WT, a slightly higher abundance of contacts between D3 and the A-helix was observed in cluster 4 for the DN mutant. This is a distinct feature, considering that these contacts are already lost in cluster 4 for the WT. **Figure 6B** shows that the interleukin tends to specifically lose contacts between D3 and helix A in the earlier stage of the reference dissociation pathway. Since the reference pathway is also present in the DN mutant, it seemed contradictory that faint bands in the contact map occurred between D3 and the A-helix in cluster 4. However, visual inspection of all RAMD trajectories for the DN and A11 mutants revealed a few trajectories in which the exact opposite pathway to the reference pathway was observed. In these trajectories, the interleukin first lost contacts in D1 and then dissociated via D2/D3 of IL-13Rα1, as depicted in **Figure 6F** and **Supplementary Movie M4**. The first frame shows that the bound state of both proteins has a large interface with numerous contacts. The second frame shows a loosely bound state that, in contrast to the reference dissociation pathway, has lost contacts between D1 and L_AB_/L_CD_, while maintaining contacts between D2 and D3 and the A- and D-helices. The third frame shows a near dissociated state, in which the remaining contacts of the second frame are lost. The last frame shows the further dissociation of the mutant IL-13 from the receptor.

The corresponding cluster representations and contact maps for all IL-13 variants in complex with IL-13Rα1 are shown in **Figures S12** and **S13**, respectively. Like DN and A11, the contact maps of the A7 and A8 IL-13 mutants also display faint bands corresponding to contacts between D3 and the A-helix, indicating that the less populated alternative pathway is present in these two systems too (**Figure S13**).

Hierarchical clustering of the ends of the dissociation trajectories supports the observation that for most of the studied systems, the reference pathway is followed with dissociation via contacts between D1 and L_AB_ and L_CD_ (**Figure S14**). Among all contacts observed in the MD-IFP analysis, the most prominent contacts just prior to dissociation are between M66 of IL-13 and E106 of its receptor, between IL-13 residue 122 (K122 in WT, R122 in A8, and M122 in B4) and receptor residues T104 and E106. Even though contacts between D3 and the A-helix might be a feature of the alternative pathway, no frames with contacts between D3 and the A-helix just prior to dissociation were captured during the hierarchical clustering (**Figure S14**). Nevertheless, contacts between D2/D3 and the D-helix were captured prior to dissociation (for the I47A, A7, B4 and B6 mutants), further indicating that IL-13 can dissociate via D2 and D3.

## Discussion

The main aim of this study was to investigate the determinants of the residence time of IL-13 to its high and low affinity receptors. We therefore first computed the residence time of IL-13Rα2 and IL-13Rα1 in complex with WT IL-13 and a total of 19 IL-13 mutants using the τRAMD method and compared the results with experimental measurements. We then analyzed the dissociation mechanisms and identified two main dissociation pathways of IL-13 from each of the receptors.

### Computation of residence time

Prior studies in which τRAMD was used to compute relative residence times showed good agreement with calculated values for protein-small molecule complexes (**45, 48, 68**) and for protein-protein complexes (**47**). The random force magnitude used in these studies ranged from 4 to 16 kcal mol^-1^ Å^-1^ for protein-small molecule systems (**45, 48, 68**), while the force magnitude for protein-protein complexes was slightly higher with values of 17 and 19 kcal mol^-1^ Å^-1^ (**47**). In contrast, the force magnitude used in this work was notably higher at approximately 26 kcal mol^-1^ Å^-1^. This force magnitude was chosen to study the dissociation of IL-13 from its high affinity receptor IL-13Rα2 because the affinity was reported to be in the fM regime (**4**). To be able to compare the low and high affinity complex, the same force magnitude was used for IL-13 in complex with IL-13Rα1. Nevertheless, it is important to consider lower random force magnitudes for the low affinity system to see whether lower values influence the correlation or could better separate the residence times of the IL-13 variants. Test calculations revealed that a lower force magnitude of 23 kcal mol^-1^ Å^-1^ did not yield a better separation between IL-13 variants with the lowest (K138A) and the highest (WT) residence time with respect to the low affinity receptor than a force magnitude of 26 kcal mol^-1^ Å^-1^. Therefore, considering that by lowering the force magnitude the dissociation trajectories tend to become longer and the computational expense rises, we did not consider it appropriate to run a full set of calculations at a lower random force magnitude.

To investigate whether the relatively high standard deviations of residence times computed with the COM-COM criterion could be due to insufficient sampling with 90 RAMD trajectories per system, we carried out tests with eight replicas, each with 15 trajectories for WT, R44A and T121A IL-13 or 20 trajectories for D120A and A5 IL-13. These simulations resulted in no notable changes in calculated residence times and their standard deviations, indicating that sampling with six replicas and at least 15 trajectories gave a good level of convergence.

The relative residence times computed using τRAMD did not correlate with experimental values for the IL-13 variants in complex with IL-13Rα1 described by **Lupardus et al.** (**4**). A possible explanation for this is that the experimental residence times of the mutants in this dataset are of relatively low duration and close to each other so that τRAMD cannot resolve such small differences due to the relatively large uncertainties in the computed values compared to the magnitude of the residence times for this set of data (**Figure S1**, **Table S2**). In contrast, the calculated relative residence times for the IL-13 variants in complex with IL-13Rα2, described by **Lupardus et al.** (**4**), and the interleukin variants in complex with IL-13Rα1, described by **Moraga et al.** (**3**), showed good correlations with the experimental values. These two data sets include mutants with distinguishable residence times over a greater range. Therefore, all three datasets were initially analyzed separately. However, normalizing the dataset from **Moraga et al.** (**3**) to the IL-13/IL-13Rα1 dataset of **Lupardus et al.** (**4**) and displaying all datasets at once also showed a good agreement between experimental and calculated residence times.

In addition to the COM-COM stopping criterion, which is the original stopping criterion used for protein-small molecule systems (**45**), four additional criteria were introduced specifically for protein-protein complexes (**47**). The reason was that proteins usually interact via numerous contacts at specific interfaces. For the IL-13 complexes studied here, the four different criteria did not significantly alter the correlation observed except for the “by residue first” criterion. By definition, this criterion considers frames of a RAMD trajectory up to the first frame in which the average distance between binding site residues exceeds a threshold distance of 5.5 Å for the calculation of the residence time. Hence, this criterion is rather strict because it does not consider re-association of contacts which would cause the average distance between all binding site residues to fall below the threshold distance of 5.5 Å later on in the simulation, and which would, for example, be accounted for by the “by residue last” criterion. This is consistent with the fact that all residence times calculated by “the residue first” criterion are lower than the residence times calculated with all other criteria, indicating that indeed re-association of contacts during dissociation does occur. The “by residue first” criterion results in a markedly lower correlation between calculated and experimental residence times for the dataset of complexes with the low affinity receptor, IL-13Rα1 (**3**). This behavior differs from the results of **D’Arrigo et al.** (**47**) for which the system with the lowest residence times gave a better correlation with the “by residue first” criterion, possibly because it better captured early unbinding processes, whereas the other systems studied were not sensitive to the criterion used. Interestingly, the dataset derived from **Moraga et al.** (**3**) also contains complexes with low residence times: the WT has a *k*_off_-rate in the 10^-2^ s^-1^ regime, while the other mutants have *k*_off_-rates in the 10^-3^s^-1^ regime (**Figures 3E** and **3F**). We surmise that the “by residue first” criterion is highly sensitive to perturbations in loosely associated contacts, as discussed by **D’Arrigo et al.** (**47**), but that this does not necessarily lead to better estimation of residence time and may, as we see for this set of interleukin complexes lead to poorer estimation of residence time. Interestingly, the “few contacts first” criterion, which follows the same definition as the “by residue first” criterion except that the residence time is determined by the first frame in which the number of contacts falls below 50%, is not as sensitive to such perturbations.

### Dissociation pathway analysis

The second part of this study focused on the analysis of the dissociation pathways of the interleukin-receptor systems. The MD-IFP analysis provided insights into distinguishable dissociation pathways of different IL-13 variants in complex with IL-13Rα2 and IL-13Rα1 on an atomistic level. The MD-IFP results generally showed that the clusters corresponding to the bound (clusters 1-2) and loosely bound (cluster 3) states had higher populations than the metastable intermediate (cluster 4) on the dissociation trajectory. This reflects the fact that the protein complex mainly remains in the bound state during the RAMD simulations while the actual dissociation process takes place in only a small fraction of all frames. The use of a lower force magnitude during the RAMD simulations might enhance the sampling of the dissociation process itself.

By utilizing the MD-IFP analysis developed for protein-protein complexes (**47, 49**), the variations in residue-residue contacts over the course of the τRAMD dissociation trajectories were tracked and revealed two distinct dissociation pathways for interleukin from the complexes with each of its receptors. For the high affinity receptor, the reference pathway identified for WT IL-13 shows initial loss of contacts of L_AB_ and L_CD_ with D1 of the receptor and then unbinding due to loss of contacts of helices A and D with domains D2 and D3 of the receptor. In contrast, an alternative pathway was observed for IL-13 F140A, in which the opposite behavior was observed: the interleukin variant first loses the contacts of D2 and D3 with helices A and D and then unbinds through loss of contacts of L_AB_ and L_CD_ with D1. This indicates that the F140A mutation weakens the interaction between the interleukin and its receptor in site II (**3, 4, 12**), favoring unbinding through D1. Since IL-13 I47A shows a similar contact pattern, it is reasonable to argue that this mutant displays a similar dissociation behavior for the same reason. IL-13 K137A displays a dissociation pattern that seems to lie in-between WT/K138A and F140A/I47A, indicating a more even distribution of reference and alternative pathways. The observation that IL-13 F140A and I47A (and possibly K137A) show an alternative egress pathway through D1 is interesting because a disruption of these important hot-spot residues (**4**) disrupts the binding between the interleukin and site II of the receptor, thus increasing the population of the alternative pathway. For interleukin in complex with IL-13Rα1, the reference route occurs via initial loss of contacts between D2 and D3 of the receptor and the A- and D-helices of the interleukin while the longest lasting contacts are for D1 with L_AB_ and L_CD_. The reference pathway is highly populated in all the mutants. In contrast, the less populated pathway, present for mutants DN, A11, A7 and possibly A8, displays the opposite behavior where first, contacts are lost between D1 and L_AB_/L_CD_ and then the interleukin dissociates with transient contacts of its A- and D-helices with D2 and D3.

The presence of distinctly different major unbinding routes for the low and high affinity receptors suggests that it is possible to selectively target the interaction with one of the receptors while having little effect on the residence time of the complex with the other receptor. The design of small molecules or proteins to modulate IL-13 activation could therefore target the interaction with D3 or D1 to alter the balance in residence times to the low and high affinity receptors. *K_D_* and *k*_off_ are well correlated across all the mutations considered. Consequently, the observed variations in *k*_off_ are expected to be primarily driven by changes in the binding free energy of the complex rather than alterations in the dissociation pathway or the transition state for unbinding. This means that the time required to disrupt the initial key contacts is more critical from a computational perspective than exhaustive sampling of the full dissociation pathways.

Five clusters were chosen to represent the IFP content of the analyzed frames of the dissociation trajectories based on the elbow and silhouette analysis for all systems. Since the number of clusters chosen is a direct tradeoff between the complexity of the analysis and the resolution of the IFP content (**49**), five clusters might not provide sufficient resolution to capture alternative pathways with low populations. For the high affinity complexes, there is a considerable population of both the reference and alternative pathways for IL-13 F140A. Yet a clear distinction between the two pathways cannot be made using the corresponding cluster representation but rather with the IFP content of the final hierarchically clustered frames. For the low affinity complexes, the reference pathway is present in all studied systems with a high population. The alternative pathway is, however, only present with a rather low population in some systems and therefore not visible in the cluster representation but is visible with the IFP content of the final hierarchically clustered frames.

## Conclusions

In conclusion, the present study demonstrates that *τ*RAMD can capture the trends in residence time for various IL-13 variants in complex with IL-13Rα1 and IL-13Rα2. This study further shows that the application of the MD-IFP tool to the RAMD trajectories revealed differences in dissociation routes and important residue-residue contacts between IL-13 and its low and high affinity receptors that could be exploited in the design of molecular modulators of IL-13:receptor interactions with desired residence time profiles. Moreover, similar receptor dissociation mechanisms can be expected for other 4-helix bundle interleukins, many of which bind to their receptors in similar arrangements. We thus anticipate that the combination of *τ*RAMD and MD-IFP analysis can be of value in the design of cytokine mimics or inhibitors with shorter (**69**) or longer (**70**) residence receptor times.

## Author contributions

NH: Investigation, Formal analysis, Validation, Data Curation, Visualization, Writing – original draft, Writing – review & editing; MB: Methodology, Validation; GDA: Methodology, Investigation, Formal analysis, Software; DKB: Conceptualization, Methodology, Software; RCW: Conceptualization, Supervision, Funding acquisition, Writing – review & editing.

## Acknowledgements

This work was supported by the European Union’s Horizon 2020 Framework Programme for Research and Innovation under Grant Agreement 945539 (Human Brain Project SGA3) and the European Union’s Research and Innovation Program Horizon Europe under Grant Agreement No. 101147319 (EBRAINS 2.0) and by the Klaus Tschira Foundation (RCW). The provision of JUSUF computing resources at Forschungszentrum Jülich (EU Horizon Project ICEI-HBP-2020-0009) is gratefully acknowledged.

We thank Bernd Doser for the RAMD implementation in Gromacs and Stefan Richter for technical support.

## Supporting material

Supplementary Tables S1 and S2, Figures S1-S14, and Movies S1-S4

## Data Availability

Input data (coordinates and parameter files) for the RAMD simulations, representative RAMD trajectories and data for computing residence times, computed interaction fingerprints, and Jupyter Notebooks for reproducing the analysis are available at Zenodo (**71**) (https://doi.org/10.5281/zenodo.18963151). All other data are available from the corresponding author on reasonable request.

## Code and software availability

SWISS-MODEL (https://swissmodel.expasy.org/) and Maestro v. 2021-4 and 2025-1 (https://www.schrodinger.com/platform/products/maestro/) were used to prepare the structure models and are freely available for academic users.

All software used for simulations is freely available: ParmEd (https://github.com/ParmEd/ParmEd), GROMACS version 2020.5 (www.gromacs.org), GROMACS-RAMD version 2020.5-2.0 (https://github.com/HITS-MCM)). AMBER 18, 20 and 22 are available with an academic license. AMBER 24 is freely available for academic, non-profit, and government usage.

Code scripts (*τ*RAMD and MD-IFP) and Jupyter notebooks used for data processing, analysis, and plotting are available at Zenodo (**69**) (https://doi.org/10.5281/zenodo.18963151) and the MD-IFP scripts (Protein-protein version 1.1) are also available at https://github.com/HITS-MCM. Codes are written in Python v. 3.x and tested on Python v. 3.8.5.

PyMol v. 3 (https://www.pymol.org/) was used for molecular visualization and figure-making and is available with an academic or commercial license. VMD v. 1.9.4 (http://www.ks.uiuc.edu/Research/vmd/) was used to generate movies of representative trajectories and is freely available for academic, educational and research purposes.

