## Supporting Material for "On the determinants of residence times and dissociation mechanisms of complexes of interleukin-13 with its low and high affinity receptors"

**Contents**


Tables S1, S2 p. 2-3

Figures S1 – S14 p. 4-15

**Table S1: Experimentally determined $k_{off}$-rates, residence times and equilibrium dissociation constants for all IL-13 variants studied.**

| IL-13 Systems | | | | Experimental data | | |
|---|---|---|---|---|---|---|
| Numbering: Publication | | | Numbering: UniProt | $k_{\text{off}}$ [$s^{-1}$]* | $\tau$ [s] | $K_D$ [M]* |
| Lupardus et al. (4) | + IL-13R$\alpha$2 | WT | WT | $1.93 \times 10^{-6}$ | 518135 | $1.58 \times 10^{-14}$ |
| | | F107A | F140A | $9.45 \times 10^{-4}$ | 1058 | $8.91 \times 10^{-11}$ |
| | | I14A | I47A | $1.52 \times 10^{-4}$ | 6579 | $8.47 \times 10^{-12}$ |
| | | K104A | K137A | $2.71 \times 10^{-3}$ | 369 | $2.89 \times 10^{-10}$ |
| | | K105A | K138A | $1.06 \times 10^{-5}$ | 94340 | $1.16 \times 10^{-12}$ |
| | + IL-13R$\alpha$1 | WT | WT | $3.97 \times 10^{-3}$ | 252 | $1.69 \times 10^{-9}$ |
| | | F107A | F140A | NA | NA | $>1.0 \times 10^{-7}$ |
| | | I14A | I47A | NA | NA | $3.17 \times 10^{-7}$ |
| | | K104A | K137A | NA | NA | $3.93 \times 10^{-7}$ |
| | | K105A | K138A | $7.71 \times 10^{-3}$ | 130 | $5.89 \times 10^{-9}$ |
| | | R11A | R44A | $4.87 \times 10^{-3}$ | 205 | $2.60 \times 10^{-9}$ |
| | | E15A | E48A | $4.45 \times 10^{-3}$ | 225 | $1.58 \times 10^{-9}$ |
| | | V18A | V51A | $4.43 \times 10^{-3}$ | 226 | $1.36 \times 10^{-9}$ |
| | | D87A | D120A | $7.51 \times 10^{-3}$ | 133 | $2.74 \times 10^{-9}$ |
| | | T88A | T121A | $4.88 \times 10^{-3}$ | 205 | $1.75 \times 10^{-9}$ |
| | | L101A | L134A | $4.63 \times 10^{-3}$ | 216 | $1.31 \times 10^{-9}$ |
| Moraga et al. (3) | + IL-13R$\alpha$1 | WT | WT | $2.20 \times 10^{-2}$ | 45 | $4.38 \times 10^{-9}$ |
| | | A5 | A5 | $8.20 \times 10^{-3}$ | 122 | $2.94 \times 10^{-9}$ |
| | | A6 | A6 | $3.30 \times 10^{-3}$ | 303 | $1.90 \times 10^{-9}$ |
| | | A11 | A11 | $1.30 \times 10^{-3}$ | 769 | $8.40 \times 10^{-11}$ |
| | | B2 | B2 | $1.70 \times 10^{-3}$ | 588 | $2.75 \times 10^{-10}$ |
| | | A7 | A7 | $3.40 \times 10^{-3}$ | 294 | $1.18 \times 10^{-9}$ |
| | | A8 | A8 | $3.30 \times 10^{-3}$ | 303 | $2.06 \times 10^{-9}$ |
| | | B4 | B4 | $2.40 \times 10^{-2}$ | 42 | $18.00 \times 10^{-9}$ |
| | | B6 | B6 | $8.20 \times 10^{-3}$ | 122 | $5.90 \times 10^{-9}$ |
| | | DN | DN | $1.30 \times 10^{-3}$ | 769 | $8.40 \times 10^{-11}$ |

* No standard deviations or error estimates are available for the measured values.

NA: For IL-13 F140A, I47A and K137A mutants in complex with IL-13Rα1, no $k_{off}$**-rates** are available **(4)**.

**Table S2: Calculated RAMD residence times for all IL-13 variants and all criteria.**

| Systems | | | Calculated residence times [ns] | | | | |
|---|---|---|---|---|---|---|---|
| | IL-13 | | ΔCOM | By residue first | By residue last | Few contacts first | Many contacts last |
| Lupardus et al. (4) | + IL-13Rα2 | WT | 20.50 ± 1.48 | 19.02 ± 0.95 | 20.18 ± 1.50 | 20.26 ± 1.49 | 20.27 ± 1.49 |
| | | F140A | 10.38 ± 3.03 | 6.84 ± 2.85 | 10.04 ± 3.07 | 10.12 ± 3.04 | 10.18 ± 3.03 |
| | | I47A | 11.03 ± 1.36 | 7.33 ± 0.72 | 10.45 ± 1.30 | 10.54 ± 1.32 | 10.60 ± 1.30 |
| | | K137A | 9.60 ± 2.56 | 7.56 ± 1.90 | 9.22 ± 2.52 | 9.09 ± 2.42 | 9.37 ± 2.56 |
| | | K138A | 12.48 ± 5.16 | 9.90 ± 4.30 | 12.11 ± 5.11 | 12.19 ± 5.10 | 12.24 ± 5.12 |
| | + IL-13Rα1 | WT | 6.85 ± 3.20 | 4.63 ± 2.24 | 6.28 ± 3.20 | 6.54 ± 3.27 | 6.56 ± 3.25 |
| | | F140A | 8.42 ± 2.08 | 4.57 ± 2.41 | 7.77 ± 2.13 | 8.15 ± 2.12 | 8.18 ± 2.10 |
| | | I47A | 2.73 ± 0.54 | 0.36 ± 0.33 | 1.89 ± 0.82 | 2.47 ± 0.53 | 2.49 ± 0.53 |
| | | K137A | 7.10 ± 1.51 | 4.04 ± 1.57 | 6.56 ± 1.43 | 6.83 ± 1.47 | 6.86 ± 1.46 |
| | | K138A | 6.95 ± 1.95 | 3.53 ± 1.92 | 6.40 ± 1.88 | 6.67 ± 1.88 | 6.70 ± 1.88 |
| | | R44A | 3.48 ± 1.38 | 2.19 ± 1.39 | 3.04 ± 1.36 | 3.27 ± 1.37 | 3.31 ± 1.37 |
| | | E48A | 7.53 ± 1.79 | 5.08 ± 1.47 | 6.88 ± 1.79 | 7.23 ± 1.79 | 7.27 ± 1.78 |
| | | V51A | 3.95 ± 0.93 | 1.95 ± 0.77 | 3.54 ± 0.93 | 3.71 ± 0.91 | 3.77 ± 0.89 |
| | | D120A | 7.55 ± 3.39 | 4.30 ± 1.38 | 6.98 ± 3.41 | 7.21 ± 3.40 | 7.27 ± 3.39 |
| | | T121A | 5.72 ± 2.47 | 3.13 ± 2.02 | 5.17 ± 2.42 | 5.49 ± 2.48 | 5.50 ± 2.48 |
| | | L134A | 6.17 ± 1.32 | 3.13 ± 1.21 | 5.63 ± 1.24 | 5.89 ± 1.24 | 5.92 ± 1.24 |
| Moraga et al. (3) | + IL-13Rα1 | WT | 6.85 ± 3.20 | 4.63 ± 2.24 | 6.28 ± 3.20 | 6.54 ± 3.27 | 6.56 ± 3.25 |
| | | A5 | 7.48 ± 3.28 | 2.57 ± 1.31 | 6.76 ± 3.27 | 7.26 ± 3.26 | 7.28 ± 3.26 |
| | | A6 | 11.82 ± 3.40 | 2.90 ± 1.63 | 10.97 ± 3.27 | 11.52 ± 3.30 | 11.53 ± 3.31 |
| | | A11 | 13.53 ± 3.22 | 6.56 ± 3.20 | 12.97 ± 3.08 | 13.22 ± 3.12 | 13.29 ± 3.15 |
| | | B2 | 12.33 ± 3.89 | 8.60 ± 4.00 | 11.86 ± 3.96 | 12.14 ± 3.88 | 12.16 ± 3.87 |
| | | A7 | 12.67 ± 1.77 | 5.35 ± 1.56 | 12.12 ± 1.79 | 12.41 ± 1.80 | 12.42 ± 1.80 |
| | | A8 | 13.85 ± 2.78 | 3.37 ± 2.13 | 13.02 ± 2.74 | 13.49 ± 2.82 | 13.55 ± 2.81 |
| | | B4 | 11.30 ± 3.66 | 6.12 ± 3.70 | 10.70 ± 3.70 | 11.07 ± 3.66 | 11.08 ± 3.67 |
| | | B6 | 10.78 ± 3.26 | 4.43 ± 2.49 | 10.12 ± 3.36 | 10.55 ± 3.24 | 10.56 ± 3.23 |
| | | DN | 19.07 ± 1.73 | 8.28 ± 4.01 | 18.37 ± 1.61 | 18.78 ± 1.73 | 18.80 ± 1.72 |

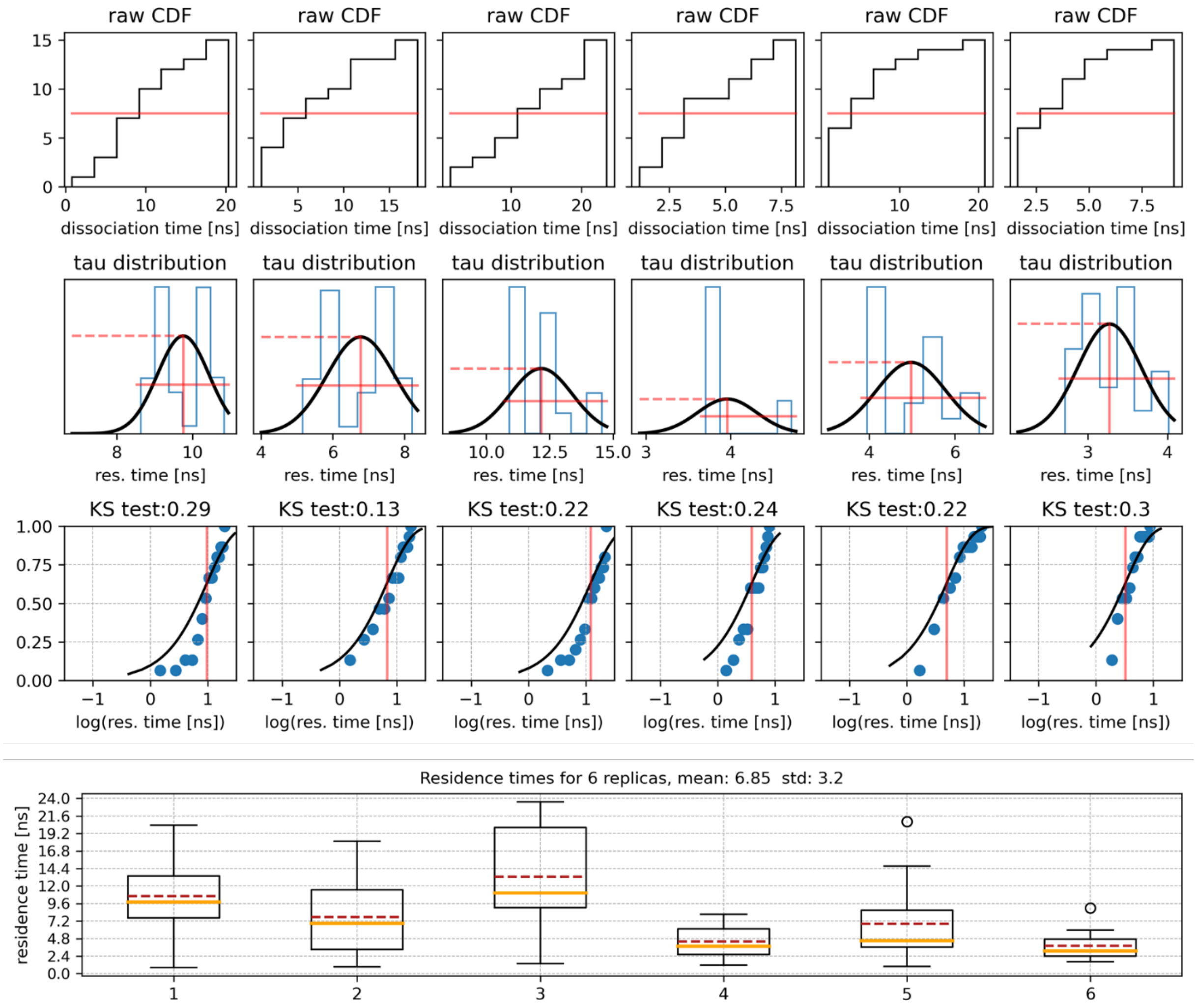


**Figure S1: Calculation and statistical evaluation of the residence time for WT IL-13 in complex with IL-13Rα1 using six replicas.**

For each of the six replicas, the cumulative distribution function is shown in panel one. The red line defines the time after which 50% of all trajectories of one replica showed a dissociation event. In the second panel, the distribution of all bootstrapped values is shown, and the third panel shows the Kolmogorov-Smirnov test to evaluate the distribution of the bootstrapped medians. The bottom panel depicts the mean residence time in red for each replica and the final calculated residence time is the average of the residence times of the six replicas **(49)**.

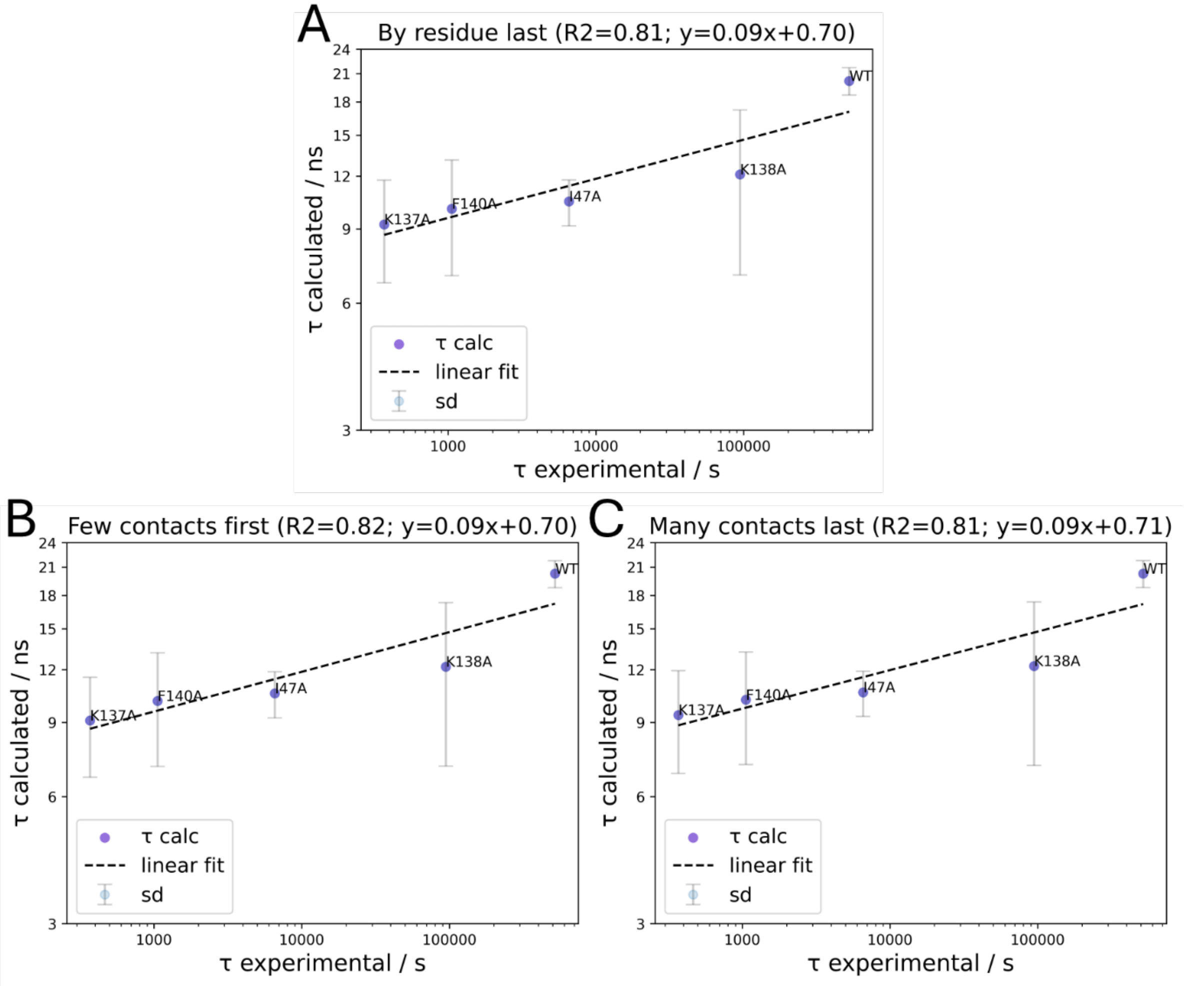


**Figure S2: Comparison of calculated and experimental residence times for WT and IL-13 mutants F140A, I47A, K137A and K138A in complex with IL-13Rα2.**

Both axes are transformed into logarithmic space. A linear regression was applied to the data points (dashed line). The data points are depicted as the mean value of six simulation replicas together with the corresponding standard deviation. The residence times were calculated according to the criteria proposed by **D'Arrigo et al. (47)**: **(A)** "by residue last" criterion, **(B)** "few contacts first" criterion, and **(C)** "many contacts last" criterion. See Fig 3. for the COM-COM and "by residue first" criteria.

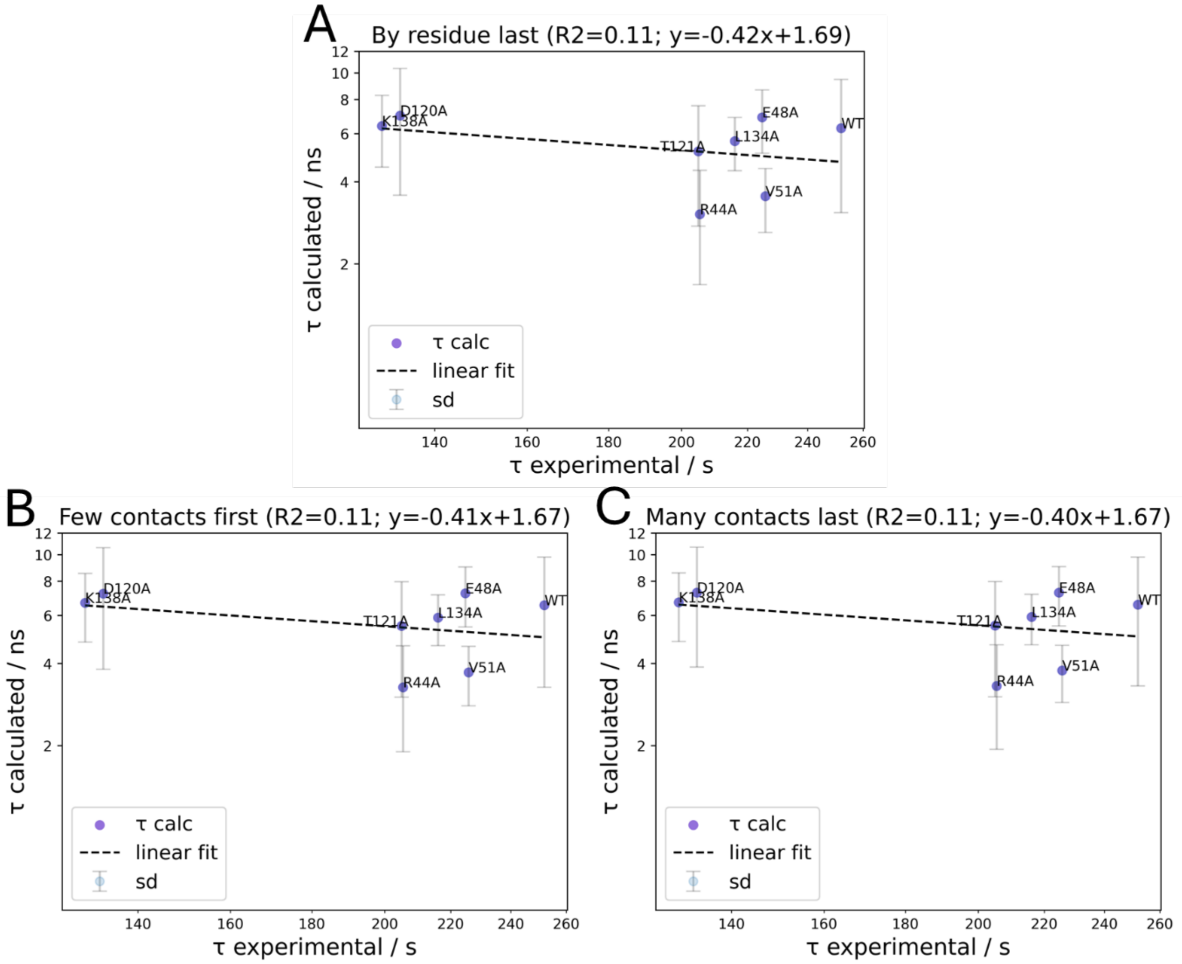


**Figure S3: Comparison of calculated and experimental residence times for WT and IL-13 mutants F140A, I47A, K137A, K138A, R44A, E48A, V51A, D120A, T121A and L134A in complex with IL-13Rα1.**

Both axes are transformed into logarithmic space. A linear regression was applied to the data points (dashed line). The data points are depicted as the mean value of six simulation replicas together with the corresponding standard deviation. The residence times were calculated according to the criteria proposed by **D'Arrigo et al. (47)**: **(A)** "by residue last" criterion, **(B)** "few contacts first" criterion, and **(C)** "many contacts last" criterion. See Fig 3. for the COM-COM and "by residue first" criteria.

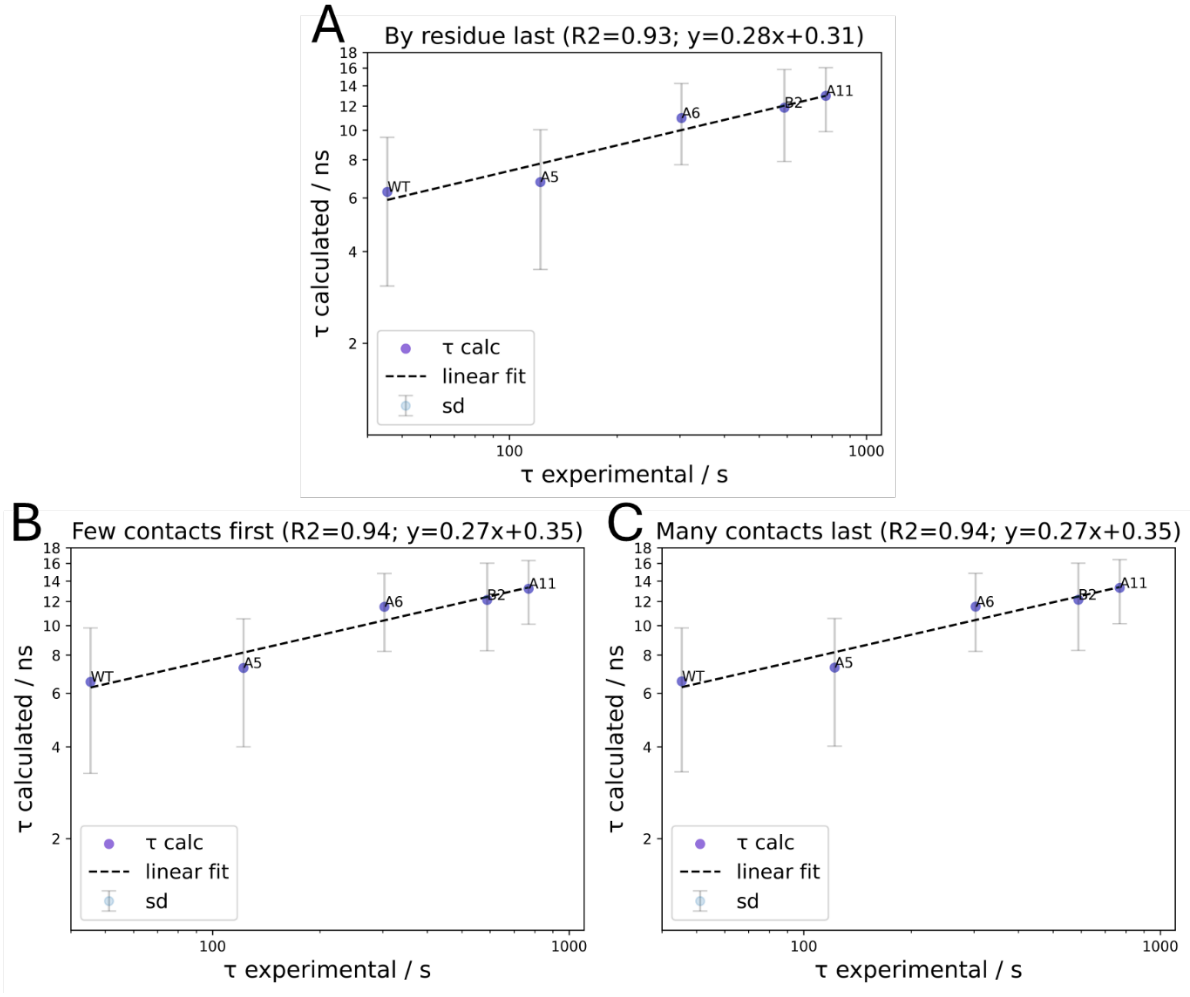


**Figure S4: Comparison of calculated and experimental residence times for WT and different IL-13 mutants A5, A6, A11 and B2 in complex with IL-13Rα1.**

Both axes are transformed into logarithmic space. A linear regression was applied to the data points (dashed line). The data points are depicted as the mean value of six simulation replicas together with the corresponding standard deviation. The residence times were calculated according to the criteria proposed by **D'Arrigo et al. (47)**: **(A)** "by residue last" criterion, **(B)** "few contacts first" criterion, and **(C)** "many contacts last" criterion. See Fig 3. for the COM-COM and "by residue first" criteria.

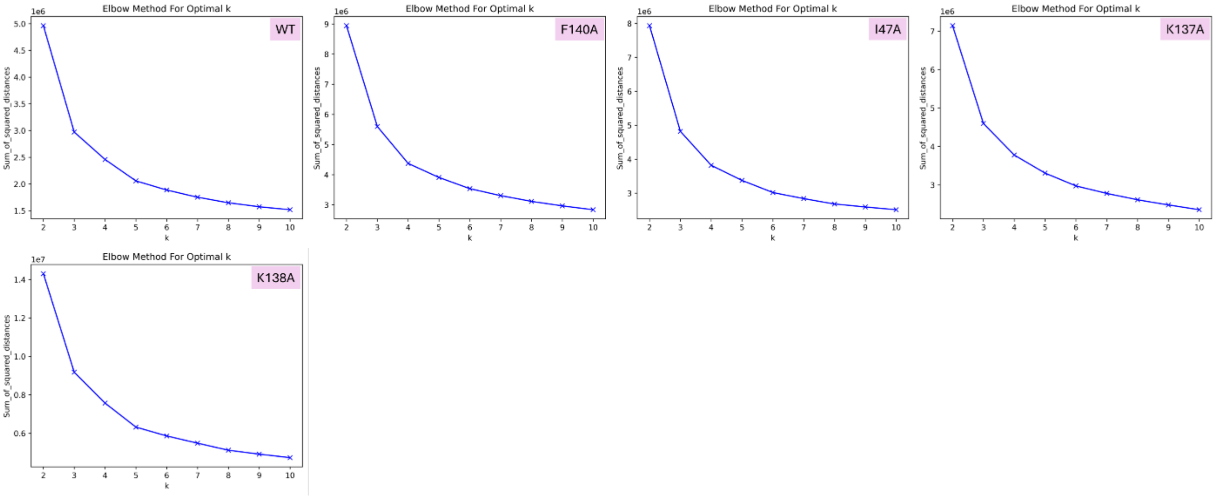


**Figure S5: Elbow plots to determine the optimal number of clusters, *k*, for MD-IFP analyses of the dissociation trajectories. Plots for all IL-13 variants studied in complexes with IL-13Rα2 are shown. 5 clusters were selected to represent most features of the dissociation routes.**

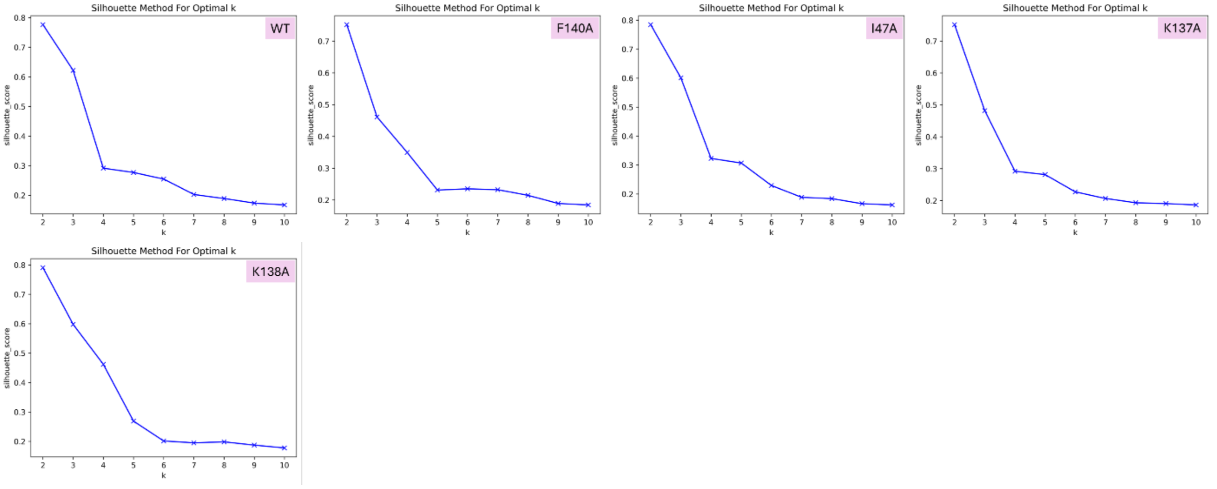


**Figure S6: Silhouette plots to determine the optimal number of clusters, *k*, for MD-IFP analyses of the dissociation trajectories for all IL-13 variants studied in complex with IL-13Rα2. 5 clusters were selected to represent most features of the dissociation routes.**

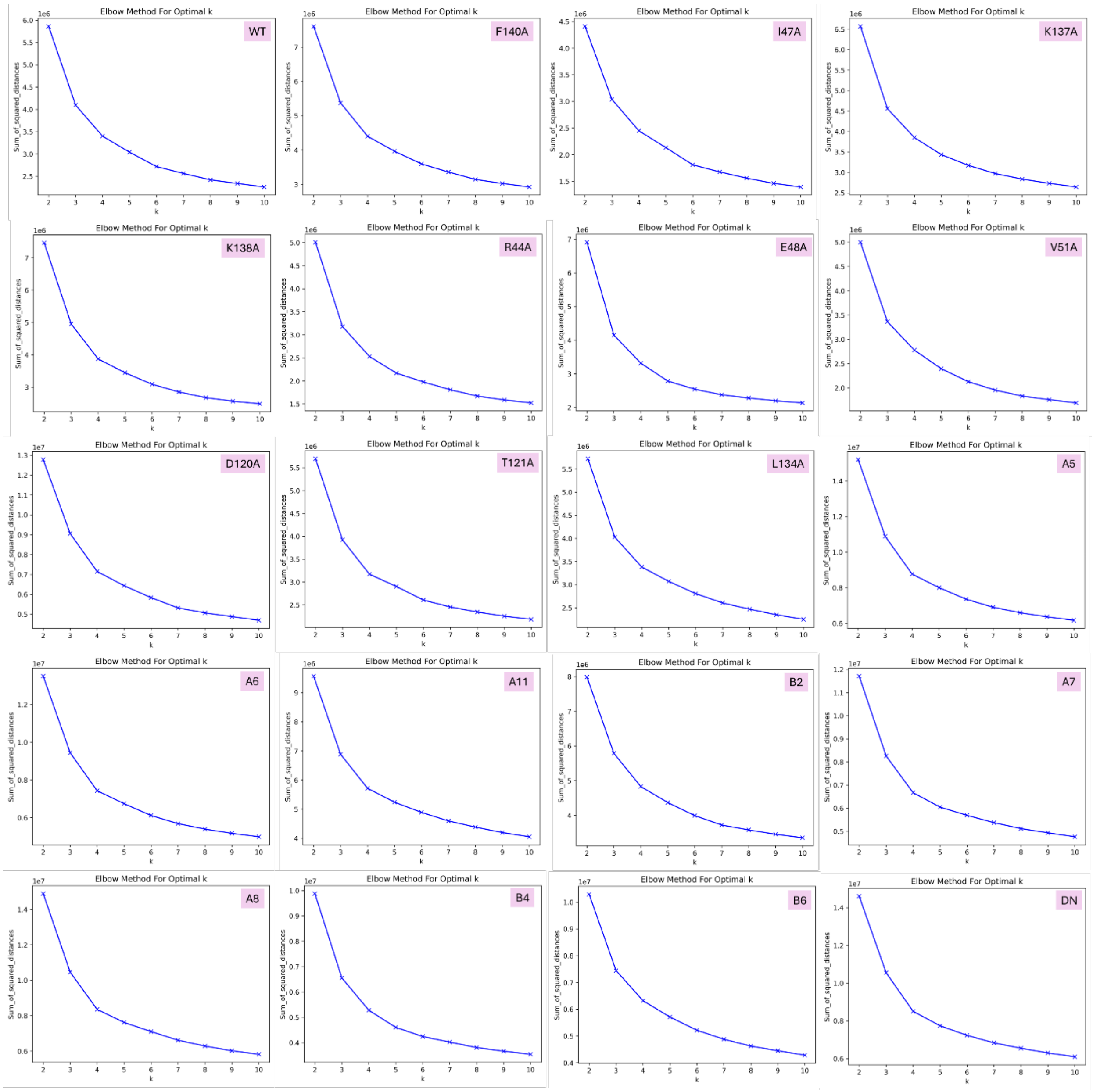


**Figure S7: Elbow plots to determine the optimal number of clusters, *k*, for MD-IFP analyses of the dissociation trajectories for all IL-13 variants studied in complex with IL-13Rα1. 5 clusters were selected to represent most features of the dissociation routes.**

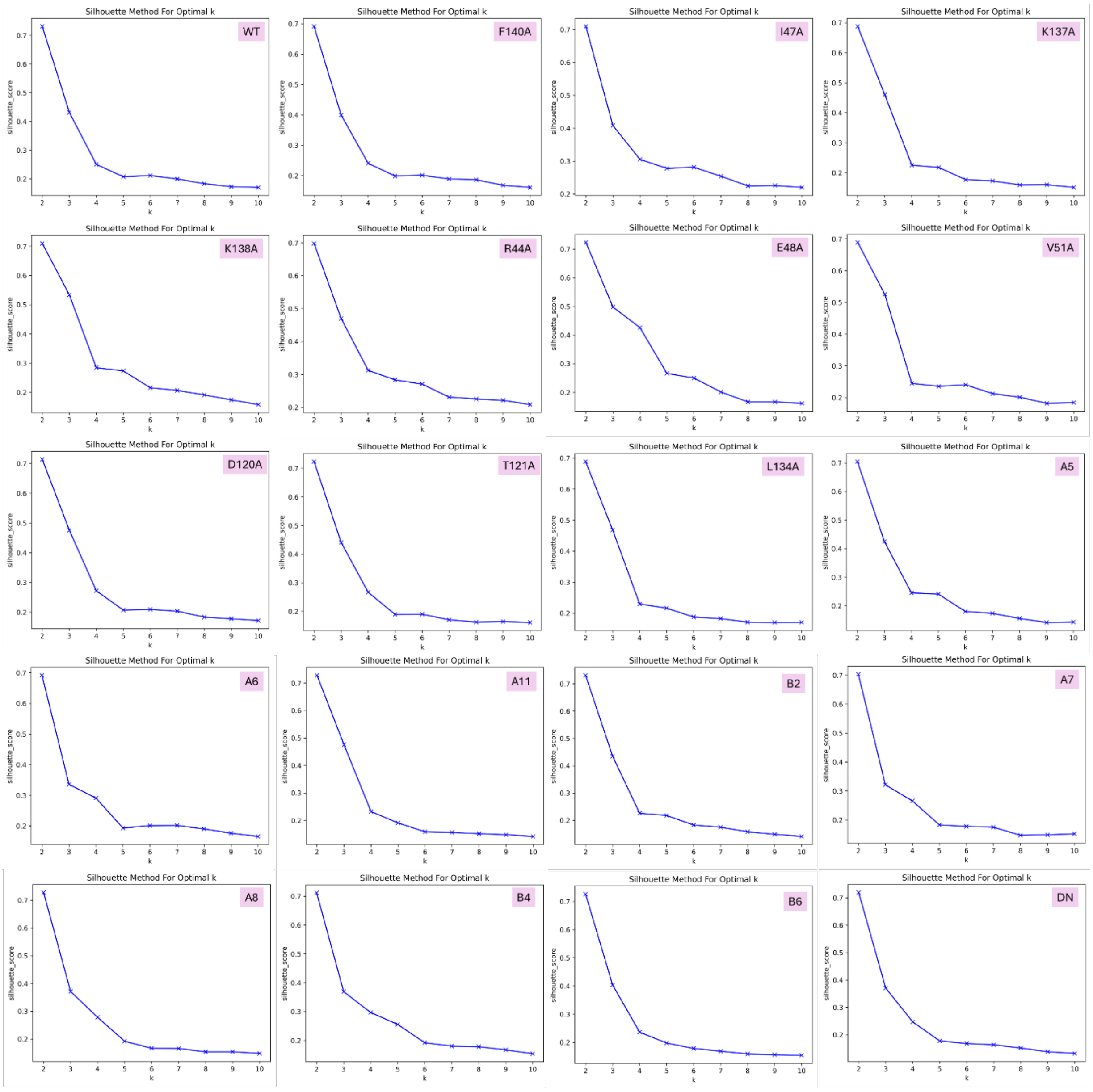


**Figure S8: Silhouette plots to determine the optimal number of clusters, *k*, for MD-IFP analyses of the dissociation trajectories for all IL-13 variants studied in complex with IL-13Rα1. 5 clusters were selected to represent most features of the dissociation routes.**

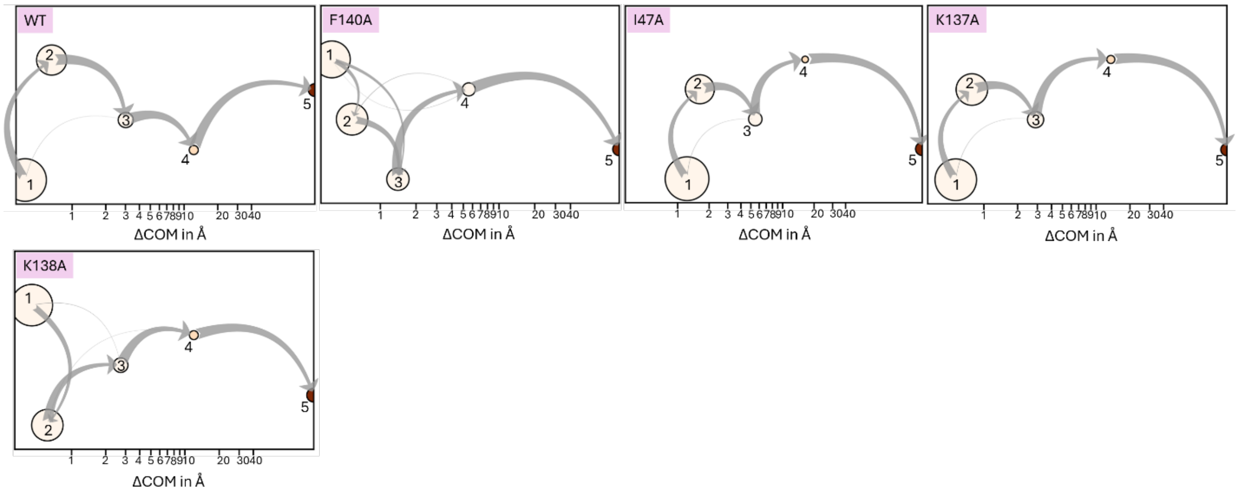


**Figure S9: Cluster representations for the dissociation trajectories of all IL-13 variants studied in complex with IL-13Rα2.**

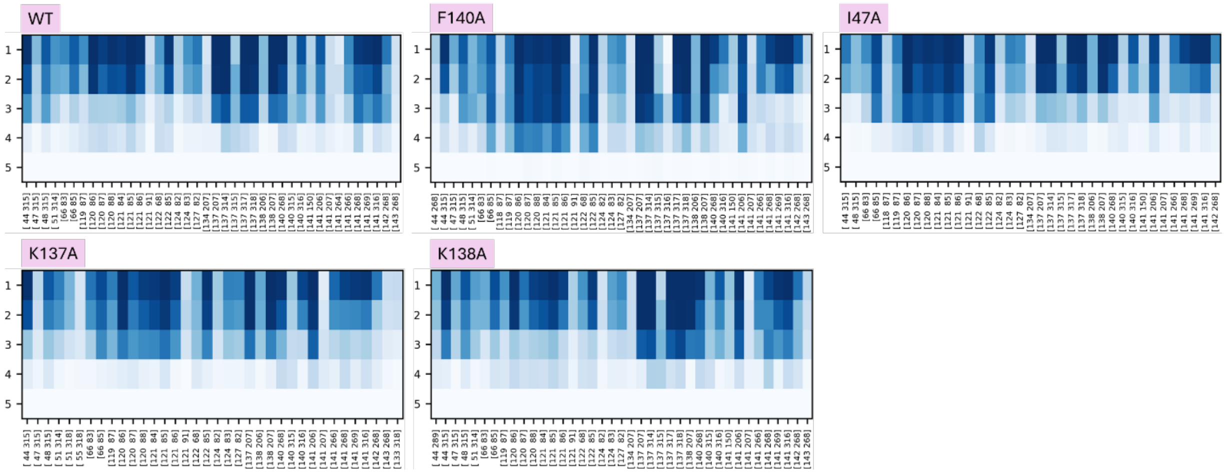


**Figure S10: Contact maps for the dissociation trajectories of all IL-13 variants studied in complex with IL-13Rα2, showing loss of residue-residue contacts on transition from the bound state (cluster 1) through different clusters to the unbound state (cluster 5).**

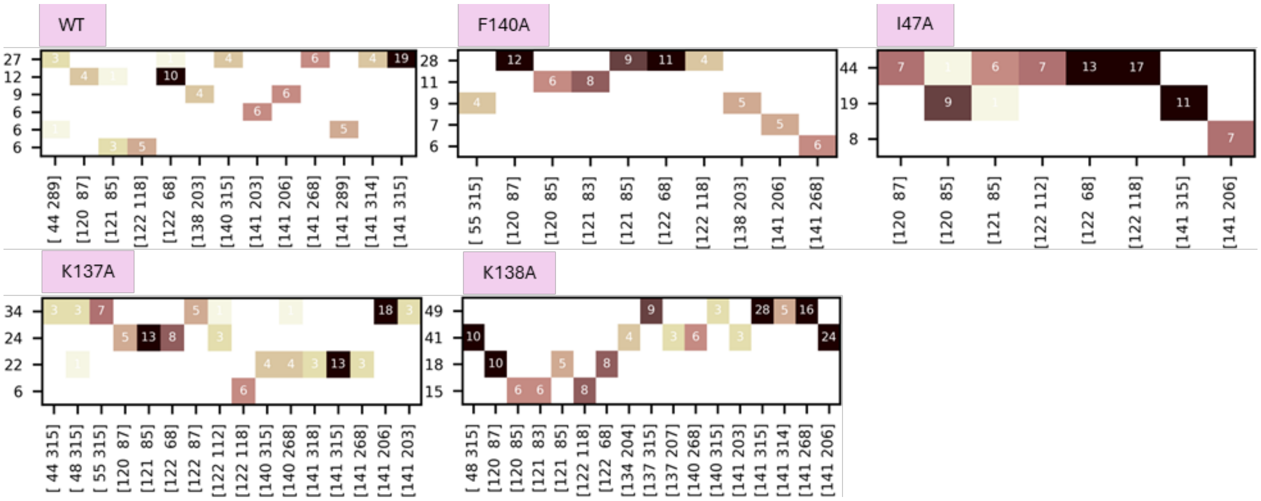

**Figure S11: Characterization of dissociation routes by hierarchical clustering of frames having fewer than three contacts towards the end of the dissociation trajectories, shown for all IL-13 variants studied in complex with IL-13Rα2.**

Each entry on the y-axis corresponds to one cluster having at least 5 frames, ordered by the number of frames. Residue-residue contacts observed in the clusters are counted (numbers in squares) and plotted on the x-axis. The higher the number, the darker the color. The residue in IL-13 is given by the lower number while the residue in the receptor is given by the upper number in the brackets.

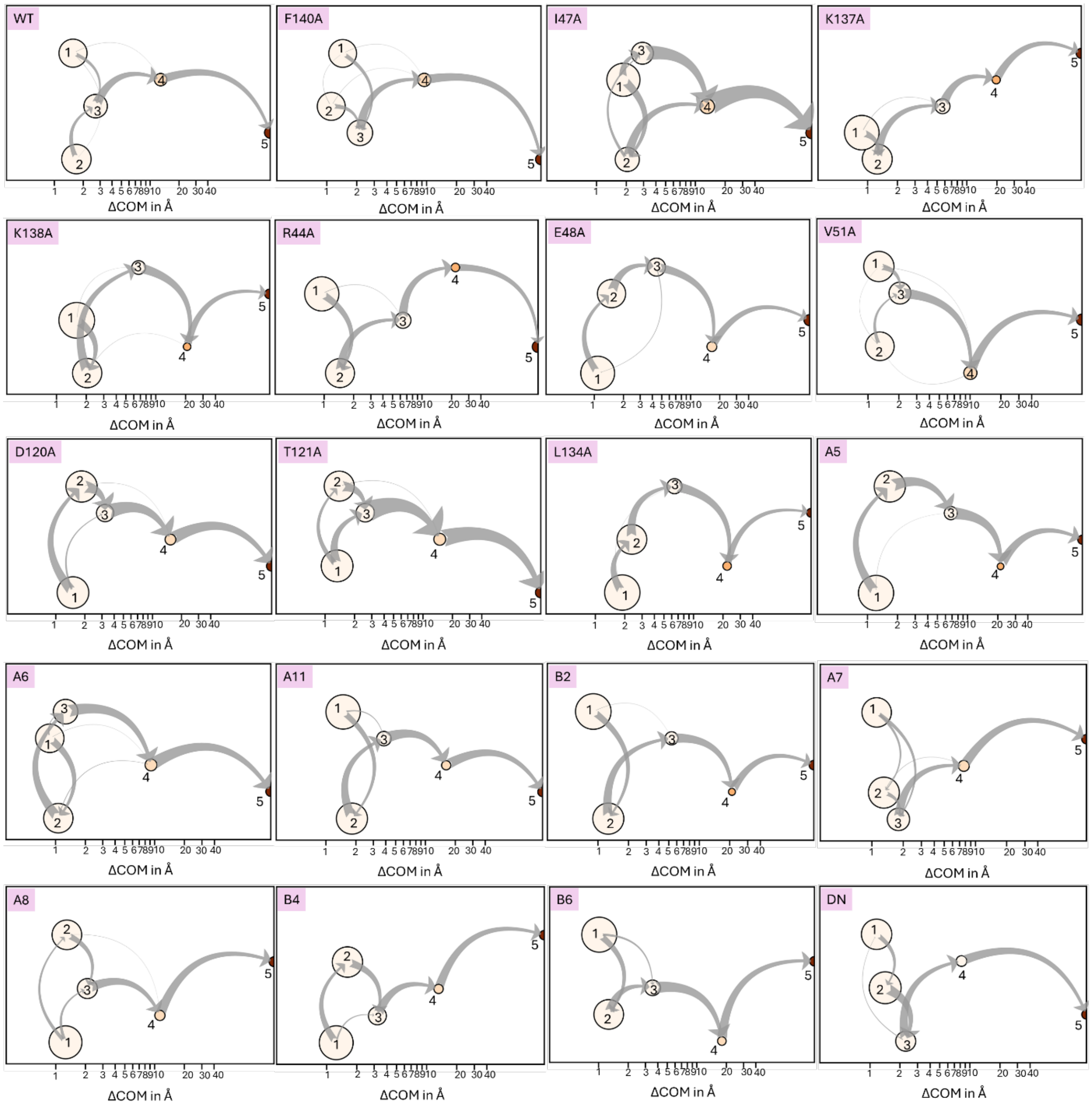


**Figure S12: Cluster representations of the dissociation trajectories for all IL-13 variants studied in complex with IL-13Rα1.**

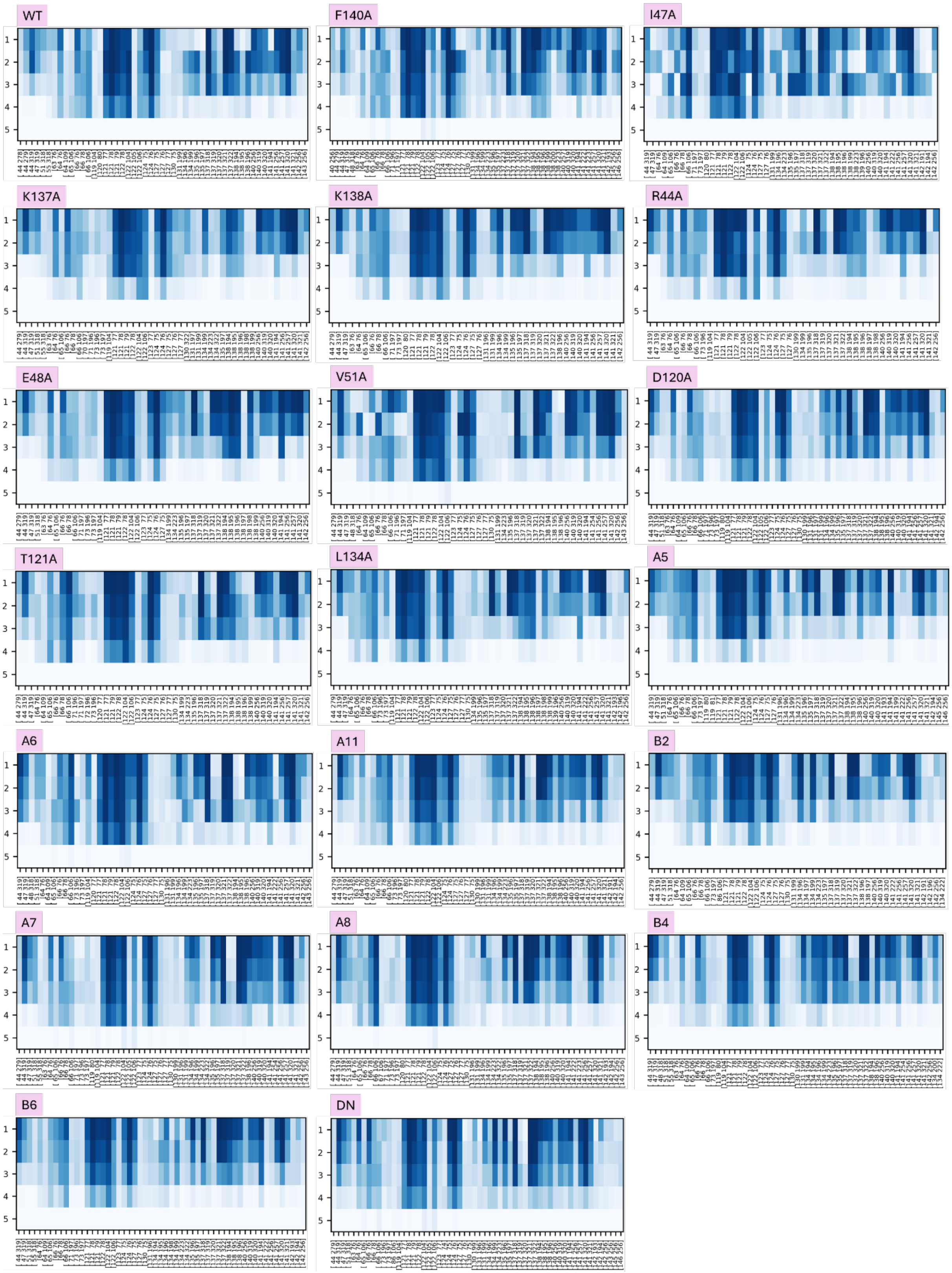


**Figure S13: Contact maps of the dissociation trajectories for all IL-13 variants studied in complex with IL-13Rα1, showing loss of residue-residue contacts on transition from the bound state (cluster 1) through different clusters to the unbound state (cluster 5).**

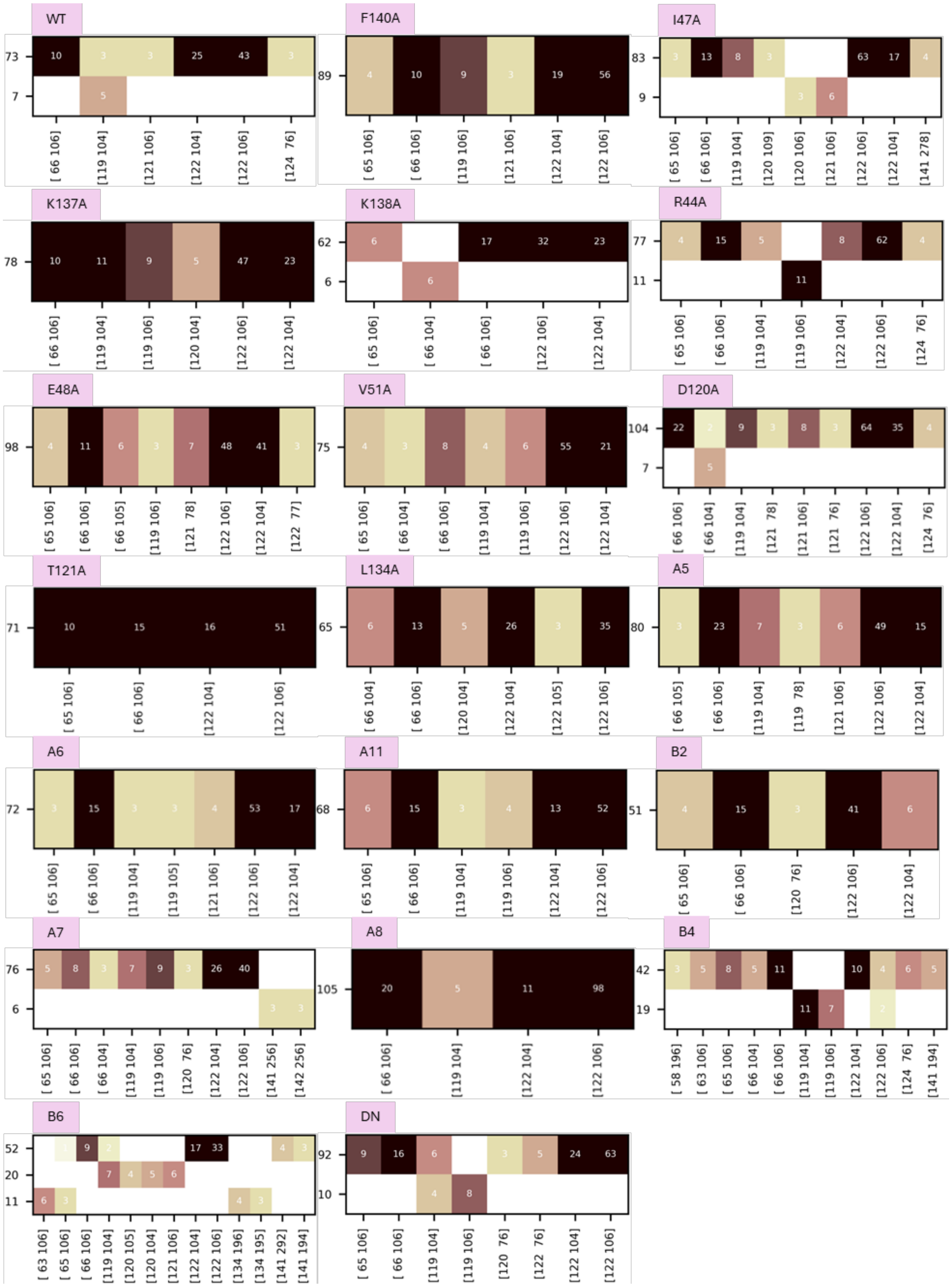

**Figure S14: Characterization of dissociation routes by hierarchical clustering of frames having fewer than three contacts towards the end of the dissociation trajectories, shown for all IL-13 variants studied in complex with IL-13Rα1. Representation as for Fig S11.**
